# Cardiomyocyte prohibitin ablation reprograms cardiac metabolism revealing a pathogenic role for mTORC1 in dilated cardiomyopathy

**DOI:** 10.64898/2026.09.01.748696

**Authors:** Ran Huo, Sarah E. Torrence, Kaitlyn A Berns, Rachel M Crawford, Amany A. Alowaisi, Jolonda C. Mahoney, Biyi Chen, Qian Shi, Benjamin W Darbro, Long-Sheng Song, Ethan J Anderson

## Abstract

Maintaining cardiac structure and function throughout the lifespan requires a delicate balance in carbon allocation between energetic and biosynthetic processes. At the nexus of this balance are prohibitins (PHB1, 2), which form a ring-like complex in mitochondrial and plasma membranes responsible for coordinating cellular growth, metabolism and autophagy. Here we describe how ablation of the PHB complex in cardiomyocytes of adult mice (cPHB1KO) causes unrestrained mechanistic target of rapamycin complex 1 (mTORC1) activity and a ‘Warburg-like’ reprogramming of glucose metabolism in heart toward enhanced *de novo* amino acid biosynthesis. These changes are accompanied by disruptions in mitochondrial Ca^2+^ handling and impaired autophagy, leading to severe dilated cardiomyopathy and mortality within 12 weeks. Using pharmacological and nutritional approaches, we further show that mTORC1 inhibition attenuates pathologic cardiac remodeling only in female cPHB1KO mice. Our findings illustrate novel mechanisms linking the PHB complex with altered carbon flux and pathogenesis of cardiomyopathy.

## Introduction

Cardiomyocytes are unique terminally differentiated cells that maintain a high rate of oxidative metabolism throughout the lifetime of an organism to sustain a massive energetic demand. For this reason, even slight alterations in carbon flux within cardiomyocytes can have substantial and long-lasting consequences to cardiac electromechanical function, illustrating why derangements in metabolism underlie almost every form of cardiomyopathy ^1–5^. A key feature of the myocardium is its ability to flexibly adapt and allocate carbon to support either ATP generation or macromolecule biosynthesis, depending on physiological needs, nutrient status and hormonal cues ^6, 7^. Our understanding of the mechanisms that disrupt cardiomyocyte metabolism and cause maladaptive cardiac hypertrophy and heart failure (HF) remains limited despite decades of research. Emerging studies have placed the prohibitin (PHB) complex squarely at the nexus of metabolic control within the cell, and in the present study we leverage the centrality and regulatory importance of this complex to interrogate metabolic mechanisms underlying cardiac hypertrophy and cardiomyopathy in detail.

PHB1 and PHB2 oligomerize to form a ringlike complex in plasma and mitochondrial membranes where they coordinate a panoply of cellular functions including growth and proliferation, metabolism and mitophagy ^8–11^. Molecular studies have demonstrated that stability of the PHB complex is dependent on both isoforms such that knockdown of either one results in destabilization of the complex and degradation of the other ^12^. Interestingly, both PHB1 and −2 have independent biological functions, though these have not been fully elucidated and are highly cell-type specific. These diverse functions of PHBs make them provocative therapeutic targets for many chronic diseases including cancer, endocrine, neurodegenerative and cardiovascular disorders ^13, 14^. However, most studies examining PHBs have been performed in proliferating cell models (e.g., cancer), making extrapolation to cardiomyocytes impossible owing to drastic differences in metabolism between these cell types. Decreased cardiac PHB1 expression has been reported in experimental HF ^15, 16^ and in HF patients ^17^, though mechanistic insights regarding the role of PHB1 in cardiomyocytes, and specifically the PHB complex, have remained elusive. Our group has recently shown that PHB1 knockdown in hepatocytes causes a decrease in AMPK activity and increase in mechanistic target of rapamycin complex 1 (mTORC1) signaling, leading to upregulation of *de novo* amino acid and lipid biosynthesis in liver ^18^.

As a highly conserved serine/threonine kinase, mTOR integrates nutrient and hormonal cues to maintain metabolic equilibrium within the cell. Like PHBs, however, most of the existing literature on mTOR is derived from work with cancer and other proliferative cell models, though some studies have examined its role in the heart (recently reviewed in ^19^). What has become quite clear from the limited studies to date is that persistent and/or excessive mTOR signaling drives cardiac hypertrophy in response to mitogen (e.g., adrenergic agonists ^20, 21^) and physiological (e.g., exercise ^22^) stimulation. A comprehensive interrogation of the molecular signals involved in mTOR-mediated cardiac hypertrophy and protein synthesis at the single cardiomyocyte level has recently been reported in an elegant study ^23^. Collectively, while these studies provide some clarity regarding mTOR signaling in the heart, the metabolic alterations that underlie cardiac hypertrophy following mTOR activation are obscure. Moreover, there are many unanswered questions concerning the mechanisms by which excessive mTOR activation becomes maladaptive and pathogenic in the heart.

To address some of these questions, in the present study we used a model of cardiomyocyte-specific PHB1 ablation (cPHB1KO) in adult mice to conditionally ablate the PHB complex in these cells. We find, surprisingly, that while PHB complex ablation has no significant effect on mitochondrial respiration in the short term, mitochondrial Ca^2+^ uptake and reactive oxygen species (ROS) are substantially affected. mTORC1 activity was also increased in the cPHB1KO hearts and only minimally responsive to fasting, while comprehensive transcript/metabolomic profiling with ^13^C-glucose revealed that cPHB1KO hearts exhibited a ‘Warburg-like’ reprogramming of glucose metabolism into *de novo* amino acid biosynthesis pathways, particularly serine and glycine synthesis. Autophagy was impaired in the cPHB1KO mice, likely owing to hyperactive mTORC1, and these mice rapidly developed severe dilated cardiomyopathy and died within 3 months of PHB1 ablation. Cardiac structural and functional remodeling occurred more rapidly in female cPHB1KO mice compared with males. Interestingly, chronic rapamycin treatment effectively blunted mTORC1 activity at the molecular level in male and female cPHB1-ko mice but was only effective at mitigating cardiomyopathy progression in females. Together these findings illustrate a role for the PHB complex in regulating mTORC1 and glucose metabolism in the myocardium and reveal a sex-dimorphic role for hyperactive mTORC1 in pathogenesis of dilated cardiomyopathy.

## Results

### Cardiomyocyte PHB complex ablation causes myocardial dilation and mortality

Cardiomyocyte-specific PHB1 knockout (cPHB1KO) mice were generated by crossing mice carrying loxP sites flanking PHB1 exon 2 with mice expressing MerCreMer recombinase under the control of the α-myosin heavy chain promoter (αMHC-MCM ^24^) (Extended Data Fig. 1). Cre-positive mice without floxxed PHB1 alleles were used as wild-type (WT) controls to account for potential Cre toxicity. Both cPHB1KO and WT mice received tamoxifen at 20 mg/kg daily for seven consecutive days, followed by serial echocardiography from 6 to 12 weeks after tamoxifen treatment (Fig. 1a). While tamoxifen had no effect in WT, survival of both male and female cPHB1KO mice began to decline approximately 10 weeks after tamoxifen treatment, and female cPHB1KO mice died approximately 1 week earlier than males (P < 0.01; Fig. 1b). Histology revealed that cPHB1KO hearts exhibited classic dilated cardiomyopathy (Fig. 1c), and serial echocardiography showed that ejection fraction (EF) was comparable between WT and cPHB1KO mice until 6 weeks after Cre induction but declined significantly thereafter (Fig. 1d). Left ventricular end-diastolic volume (EDV), end-systolic volume (ESV) and volume-to-mass ratio were also unchanged at 6 weeks but increased significantly thereafter in male and female cPHB1KO mice, indicative of progressive ventricular dilation. Heart weight-to-body weight ratios were unchanged at 6 weeks (Fig. 1e) but were significantly higher in cPHB1KO mice than in WT controls at 12 weeks (Fig. 1f). RNA sequencing demonstrated a marked reduction in PHB1 transcripts in male and female cPHB1KO hearts (Fig.1g) and immunoblot confirmed depletion of PHB1 protein in adult cardiomyocytes (Fig. 1h). PHB2 protein abundance was also reduced to similar extent as PHB1, consistent with the interdependence of PHB1 and PHB2 to form a stable complex ^12^.

**Fig. 1:**
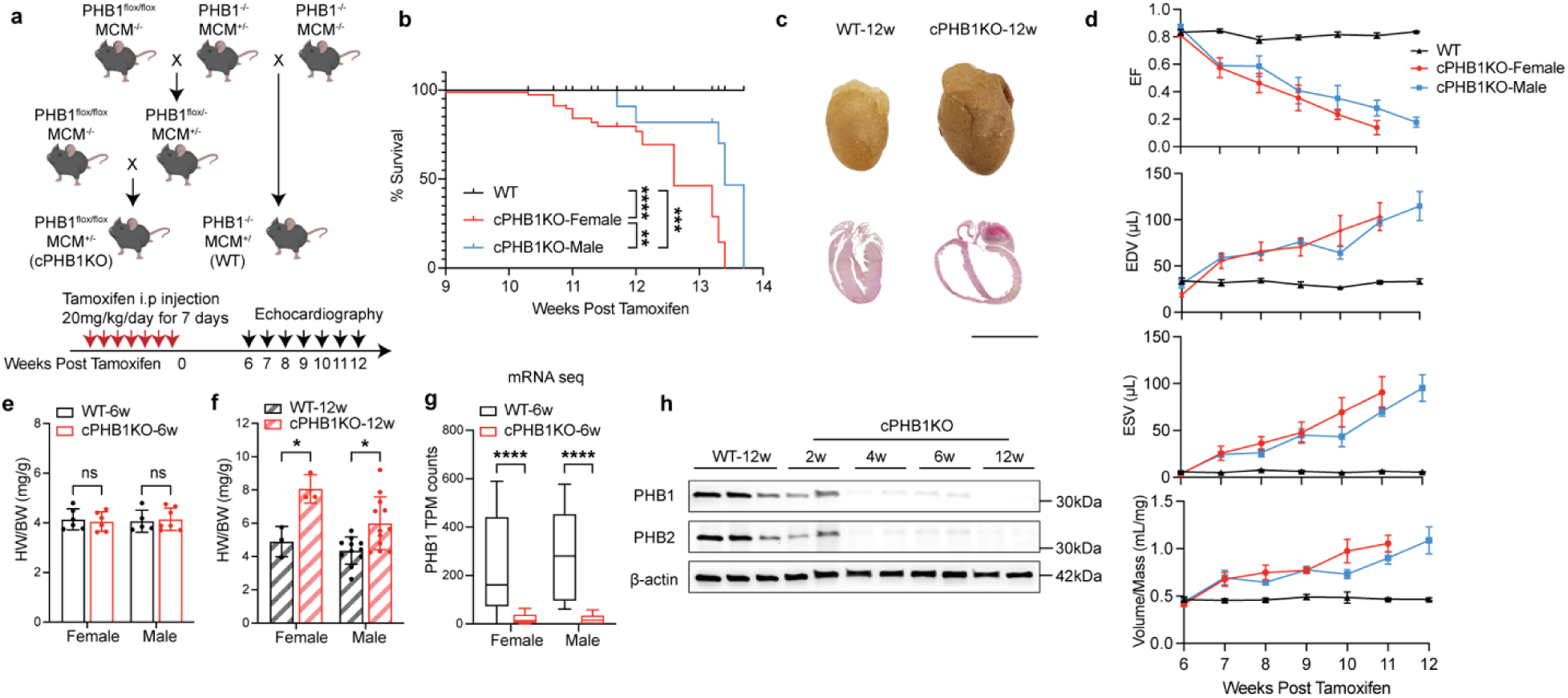
Cardiomyocyte-specific PHB1 deletion leads to concurrent loss of PHB2 and dilated cardiomyopathy. **a**, Schematic of the tamoxifen-inducible, cardiomyocyte-specific PHB1-knockout (cPHB1KO) mouse model and its wild-type (WT) control, and experimental timeline. **b,** Kaplan–Meier survival curves for WT and cPHB1KO mice (WT, *n =* 29; female cPHB1KO, *n =* 25; male cPHB1KO, *n =* 16 mice). P values were calculated by log-rank (Mantel–Cox) test. **c,** Representative images of fixed whole hearts and hematoxylin and eosin (H&E)-stained sections from WT and cPHB1KO mice at 12 weeks post-tamoxifen (WT-12w and cPHB1KO-12w). Scale bars, 5mm. **d,** Echocardiographic quantification of left ventricular ejection fraction (EF), end-diastolic volume (EDV), end-systolic volume (ESV) and volume-to-mass ratio (Vol/mass) over time. Data are presented as mean ± s.e.m. WT, *n =* 3; female cPHB1KO, *n =* 4; male cPHB1KO, *n = 4* mice. Measurements from week 6 to week 11 were analyzed by two-way repeated-measures ANOVA with Šidák’s multiple-comparisons test versus WT at each timepoint, the group × time interaction was significant for all parameters (EF and EDV, P < 0.0001, ESV and Vol/mass, P < 0.0005). Week-12 data represent surviving animals only and were not included in the primary longitudinal analysis. **e,f,** Ratio of heart weight (HW) to body weight (BW) at 6 weeks (**e**) and 12 weeks (**f**) post-tamoxifen in WT and cPHB1KO mice. Data are presented as mean ± s.d. Female: WT-6w, *n = 6*; cPHB1KO-6w, *n = 6*; WT-12w, *n = 3*; cPHB1KO-12w, *n = 3*. Male: WT-6w, *n = 5*; cPHB1KO-6w, *n = 7*; WT-12w, *n = 11*; cPHB1KO-12w, *n = 12* mice. **g,** PHB1 transcripts per million (TPM) from mRNA sequencing of hearts at 6 weeks post-tamoxifen (*n =* 6 mice per group). **h,** Representative western blot of PHB1 and PHB2 in adult cardiomyocytes isolated from WT mice at 12 weeks post-tamoxifen and from cPHB1KO mice at 2, 4, 6 and 12 weeks (2w, 4w, 6w, 12w) post-tamoxifen. β-actin was used as a loading control. For bar graphs (**e–g**), *P* values were calculated by two-way ANOVA with Tukey’s multiple-comparisons test. \**P* < 0.05, \*\**P* < 0.005, \*\*\**P* < 0.0005, \*\*\*\**P* < 0.0001, n.s., not significant.

### Mitochondrial respiration and ultrastructure in hearts are minimally affected by PHB complex ablation

Given that cPHB1KO hearts had barely detectable PHB1/2 protein but still maintained normal structure and function at 6 weeks post-tamoxifen, this represented a pre-symptomatic ‘window’ to investigate mechanisms linking PHB complex ablation to the development of dilated cardiomyopathy. At 6 weeks after Cre induction with tamoxifen, mitochondrial respiration (*J*O₂) and ATP production (*J*ATP) supported by pyruvate, palmitoyl-carnitine or glutamate were unchanged in cPHB1KO hearts of both sexes (Fig. 2a,b and Extended Data Fig. 2a,b). By 12 weeks, however, *J*O₂ was significantly reduced in both sexes across all substrate conditions (Fig. 2c), whereas *J*ATP was significantly decreased in males and showed a non-significant trend toward reduction in females (Fig. 2d). Consistent with these functional defects, OXPHOS complexes I–V were comparable between genotypes at 6 weeks but significantly reduced in cPHB1KO hearts of both sexes at 12 weeks (Fig. 2e–j). Blue native PAGE analysis showed no differences in individual respiratory complexes or supercomplexes (Fig. S2e–g) at 6 weeks. Interestingly, reverse electron transport (RET) driven H₂O₂ production was reduced in permeabilized cardiac fibers from females and isolated mitochondria from males at 6 weeks (Extended Data Fig. 2c,d). By 12 weeks, RET driven H₂O₂ production was markedly reduced in both sexes. Transmission electron microscopy of left ventricle revealed less organized mitochondrial cristae in cPHB1KO hearts at 6 weeks post tamoxifen (Fig. 2k). Despite these ultrastructural alterations, mitochondrial membrane fluidity assessed by MC540 fluorescence emission scans (540 - 620 nm) was unchanged between genotypes in either the absence or presence of BSA, with comparable peak fluorescence at 580 nm (Fig. 2l,m).

**Fig. 2:**
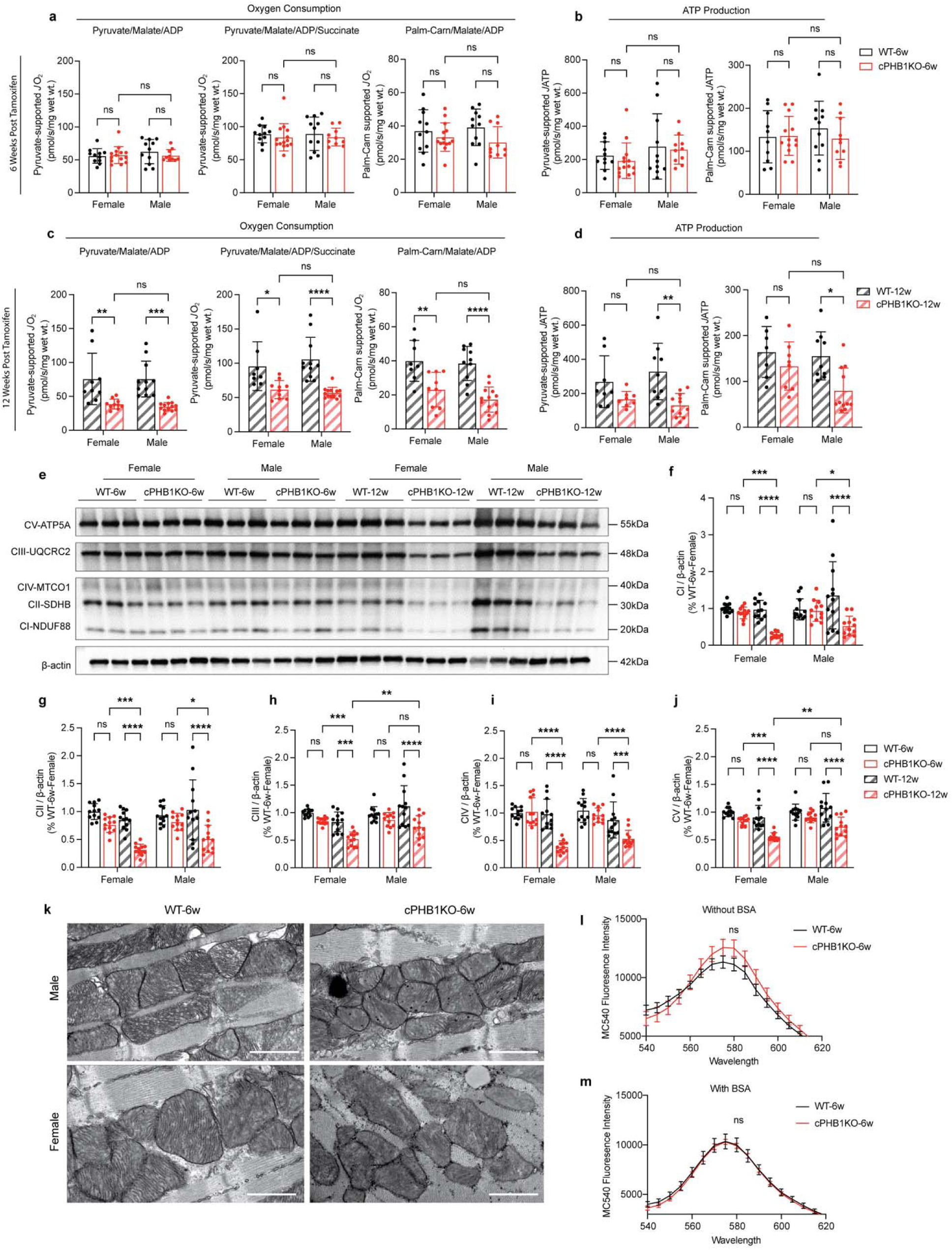
Ablation of PHB complex causes minimal changes in cardiac mitochondrial respiration and ultrastructure until late stages of heart failure. **a,b**, Mitochondrial respiratory flux (*J*O₂) (a) and ATP production (*J*ATP) (b) supported by pyruvate and palmitoyl-carnitine (Palm-Carn) in permeabilized myocardial fibers from mice at 6 weeks post-tamoxifen. Data are presented as mean ± s.d. Female: WT-6w, *n* = 11; cPHB1KO-6w, *n* = 14. Male: WT-6w, *n* = 11; cPHB1KO-6w, *n* = 10 mice. Significance was determined by two-way ANOVA with Tukey’s multiple-comparisons test. c,d, *J*O₂ (c) and *J*ATP (d) supported by pyruvate and palmitoyl-carnitine in permeabilized myocardial fibers from mice at 12 weeks post-tamoxifen. Data are presented as mean ± s.d. Female: WT-12w, *n* = 7–9; cPHB1KO-12w, *n* = 8–10. Male: WT-12w, *n* = 9–11; cPHB1KO-12w, *n* = 11–13 mice. Significance was determined by two-way ANOVA with Tukey’s multiple-comparisons test. Substrate conditions were pyruvate/malate/ADP and palmitoyl-carnitine/malate/ADP (maximal capacity), and pyruvate/malate/ADP/succinate (combined complex I and II, CI and CII). e, Representative immunoblot of mitochondrial OXPHOS complex proteins in left ventricular tissue extracts from female and male WT and cPHB1KO mice at 6 and 12 weeks post-tamoxifen. Data are representative of three biological replicates. β-actin was used as the loading control. f–j, Quantification of OXPHOS complexes I–V (CI–CV) from four immunoblots (*n* = 12 biological replicates per group). Data are presented as mean ± s.d. Significance was determined by two-way ANOVA with Tukey’s multiple-comparisons test. k, Representative transmission electron microscopy (TEM) images of cardiac mitochondria in the left ventricle from WT and cPHB1KO mice at 6 weeks post-tamoxifen. Magnification, ×20,000. Scale bar, 1 µm. **l**,**m**, Mitochondrial membrane fluidity assessed by MC540 emission spectra (540–620 nm) in the absence (**l**, *n* = 8 per group) and presence (**m**, *n* = 12 per group) of BSA. Curves show mean ± s.e.m. Peak MC540 fluorescence intensity at 580 nm was compared between genotypes by unpaired two-tailed *t*-test (WT-6w versus cPHB1KO-6w, sexes pooled), differences were not significant (**l**, *P* = 0.14). For all panels, \**P* < 0.05, \*\**P* < 0.01, \*\*\**P* < 0.001, \*\*\*\**P* < 0.0001, n.s., not significant.

### PHB complex ablation disrupts mitochondrial Ca²⁺ handling, membrane potential and ROS in cardiomyocytes

Mitochondrial Ca²⁺ uptake rate and retention capacity were determined by applying sequential 10 μM Ca²⁺ pulses to isolated mitochondria until mitochondrial permeability transition pore (mPTP) opening (Fig. 3a). At 6 weeks post-tamoxifen, Ca²⁺ retention capacity was significantly increased in female cPHB1KO mitochondria compared with WT, whereas no genotype difference was observed in males (Fig. 3b). Further analysis revealed an increased Ca²⁺ uptake rate in cPHB1KO mitochondria of both sexes, with a greater increase in females (Fig. 3c,d). Addition of recombinant PHB1, alone or together with recombinant PHB2, reduced the elevated Ca²⁺ retention capacity in female cPHB1KO mitochondria (Extended Data Fig. 3a) and attenuated the increased Ca²⁺ uptake rate in both sexes (Extended Data Fig. 3b,c,e,f) to levels similar with WT. In males, rPHB1 alone or combined with rPHB2 reduced Ca²⁺ retention capacity in both WT and cPHB1KO mitochondria (Extended Data Fig. 3d). We next examined components of the mitochondrial Ca²⁺ uniporter complex (Fig. 3e). MICU1 abundance was unchanged between genotypes at 6 weeks but markedly reduced in cPHB1KO hearts of both sexes at 12 weeks (Fig. 3f). MCU abundance was similarly unchanged at 6 weeks, whereas at 12 weeks it was significantly reduced in female cPHB1KO hearts (Fig. 3g). To gain further insights on mitochondrial organization and energetics at a macroscale, we next used *in situ* confocal imaging of tetramethylrhodamine ethyl ester (TMRE) and 2’, 7’-dichlorofluorescein (DCF) to measure mitochondrial membrane potential and ROS production, respectively, in non-beating intact hearts (Fig. 3h). Though not quantified, TMRE revealed that cPHB1KO cardiomyocytes also exhibited a less organized mitochondrial staining pattern compared with WT hearts. Analysis of TMRE fluorescence intensity showed no significant differences between genotypes within either sex at 6 weeks, although female cPHB1KO hearts had significantly lower TMRE fluorescence intensity than male cPHB1KO hearts (Fig. 3i). In contrast, DCF fluorescence was greatly increased in cPHB1KO hearts from both sexes compared with WT, with a greater increase in females (Fig. 3j). While collectively these findings support prior studies implicating the PHB complex in mitochondrial respiration, ROS and Ca²⁺ handling, they cannot fully explain pathogenesis of cardiomyopathy in the cPHB1KO hearts, suggesting additional factors are involved.

**Fig. 3:**
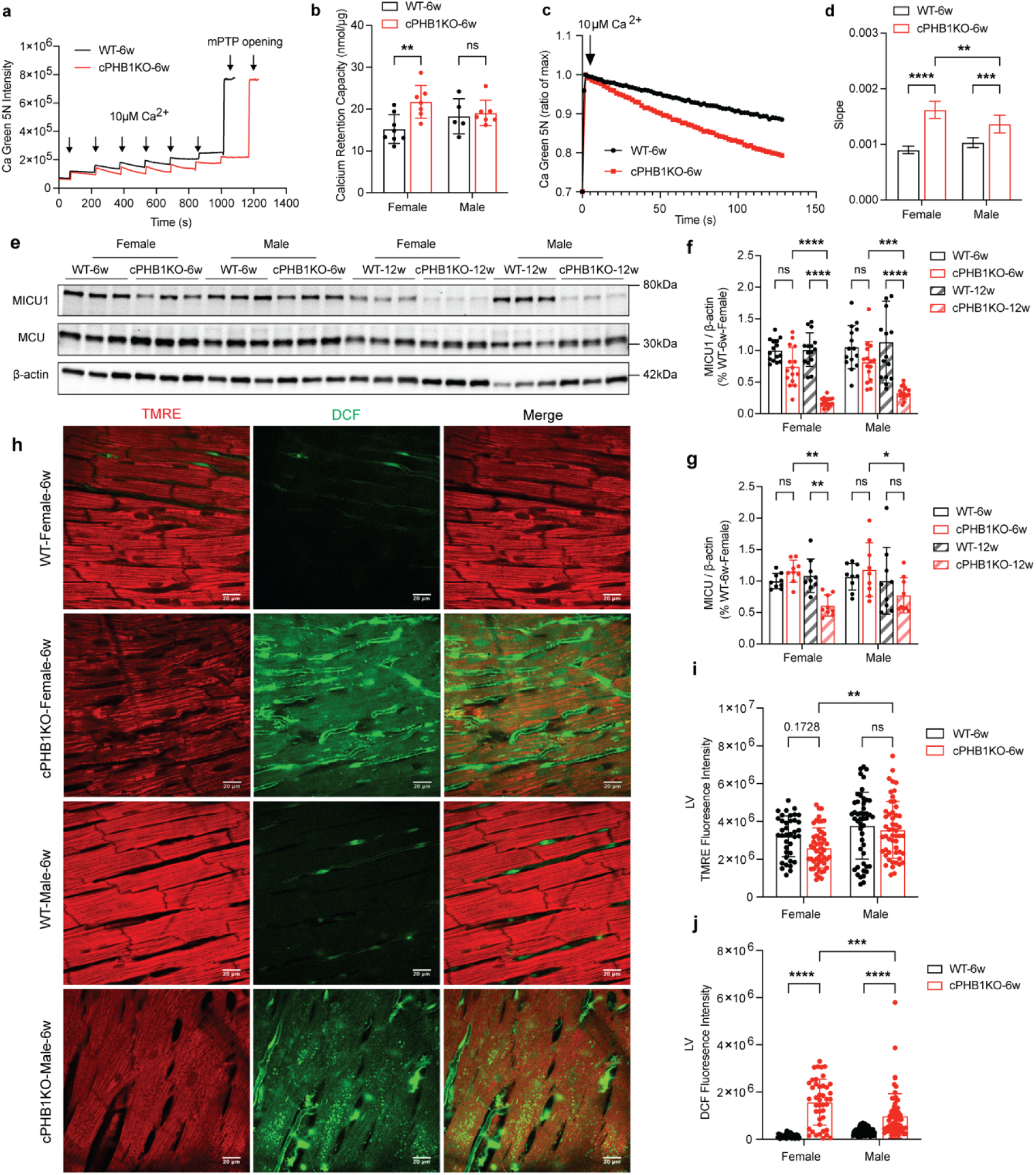
PHB complex ablation disrupts *in situ* mitochondrial Ca^2+^ handling, membrane potential and ROS production in the heart. **a**, Representative mitochondrial calcium-uptake traces from cardiac mitochondria measured with an Oroboros fluorimeter. Arrows indicate the timing of calcium pulses and mitochondrial permeability transition pore (mPTP) opening. b, Quantification of calcium retention capacity from traces as in a. Data are presented as mean ± s.d. Female: WT-6w, *n* = 8; cPHB1KO-6w, *n* = 7. Male: WT-6w, *n* = 5; cPHB1KO-6w, *n* = 7 mice. c, Representative traces of the second calcium-uptake peak from a. d, Slope of the linear regression fitted to the descending phase of the second fluorescence peak in c. Data are presented as mean ± s.d. Significance was determined by two-way ANOVA with Tukey’s multiple-comparisons test. Female: WT-6w, *n* = 8; cPHB1KO-6w, *n* = 7. Male: WT-6w, *n* = 6; cPHB1KO-6w, *n* = 7 mice. e, Representative western blot of mitochondrial calcium uniporter complex proteins in left ventricular tissue extracts from female and male WT and cPHB1KO mice at 6 and 12 weeks post-tamoxifen. Data are representative of three biological replicates. β-actin was used as the loading control. f,g, Quantification of MICU1 (f, from five immunoblots, *n* = 15 biological replicates per group) and MCU (g, from three immunoblots, *n* = 9 biological replicates per group). Data are presented as mean ± s.d. h, Representative in situ confocal images of left ventricle (LV) stained with TMRE and DCF to assess mitochondrial membrane potential (ΔΨm) and cytosolic reactive oxygen species (ROS), respectively. Scale bar, 20 μm i,j, Quantification of TMRE (i) and DCF (j) fluorescence intensity (10–20 images per mouse, *n* = 3–5 mice per group). Data are presented as mean ± s.d. For all bar graphs, statistical significance was determined by two-way ANOVA with Tukey’s multiple-comparisons test. \**P* < 0.05, \*\**P* < 0.01, \*\*\**P* < 0.001, \*\*\*\**P* < 0.0001, n.s., not significant.

### Loss of cardiomyocyte PHB complex causes mTOR hyperactivation and altered glucose and amino acid metabolism in the heart

Our group ^18^ and others ^25, 26^ ^27^ have recently found that PHBs are linked to mTORC1 signaling in multiple cell types. Thus, we comprehensively examined mTORC1 signaling in WT and cPHB1KO hearts following Cre induction by tamoxifen (Fig. 4a). Phosphorylation of 4EBP1 at Ser64 was significantly increased in cPHB1KO hearts at 6, 9 and 12 weeks, whereas total 4EBP1 was unchanged. Phosphorylation of p70S6K also appeared to increase in the cPHB1KO hearts in parallel with p-4EBP1, though the quantitative differences were masked by the increase in total p70S6K that occurred congruently in these hearts (Fig. 4b).

**Fig.4:**
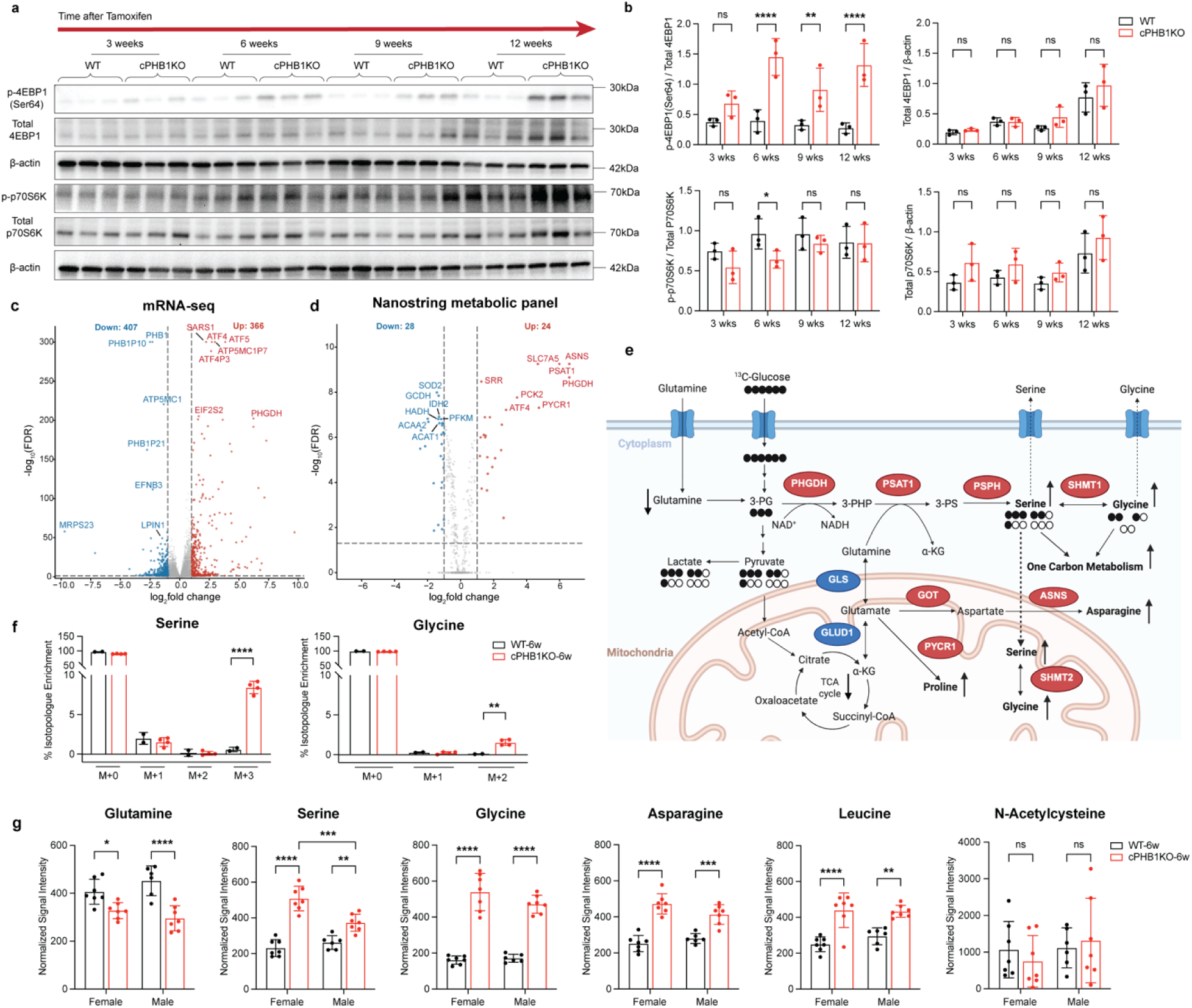
Loss of PHB complex leads to mTOR hyperactivation and reprogramming of glucose metabolism in the heart. **a**, Western blot of the mTOR-pathway proteins phospho-4EBP1 (p-4EBP1) and phospho-p70S6 kinase (p-p70S6K) at 3, 6, 9 and 12 weeks after tamoxifen in female WT and cPHB1KO mouse hearts. Gels/blots were processed in parallel. Data are representative of three biological replicates. β-actin was used as the loading control. b, Quantification of the immunoblots in a. Data are presented as mean ± s.d. Statistical significance was determined using two-way ANOVA followed by Šídák’s multiple-comparisons test. c,d, Volcano plots of differentially expressed genes in left ventricle of cPHB1KO versus WT mice at 6 weeks post-tamoxifen, from mRNA sequencing (c, *n = 12 per group (WT and cPHB1KO)*, 6 female and 6 male) and NanoString metabolic profiling (d, *n* = 6 per group *(WT and cPHB1KO)*, 3 female and 3 male). e, Schematic of glutamine metabolism and the serine/glycine biosynthesis pathway with ¹³C isotope tracing. Created with BioRender. f, Quantification of ¹³C isotope tracing. WT-6w, *n* = 2; cPHB1KO-6w, *n* = 4 mice. Data are presented as mean ± s.d. Significance was determined by unpaired two-tailed Welch’s *t*-test. g, Abundance of selected metabolites associated with the pathways shown in e. WT-6w male, *n* = 6; WT-6w female, cPHB1KO-6w female and cPHB1KO-6w male, *n* = 7 each. Data are presented as mean ± s.d. Significance was determined by two-way ANOVA with Tukey’s multiple-comparisons test. For all panels, \**P* < 0.05, \*\**P* < 0.01, \*\*\**P* < 0.001, \*\*\*\**P* < 0.0001, n.s., not significant.

To gain further insights using an orthogonal approach, we performed mRNA sequencing (Fig, 4c) and targeted NanoString profiling (Fig, 4d) in bulk cardiac tissue from the WT and cPHB1KO mice at 6 weeks following Cre induction. While canonical signatures of cardiac hypertrophy were unremarkable in the cPHB1KO hearts (Extended Data Fig. 4), there was clear up-regulation of integrated stress response (ISR) and amino acid biosynthetic genes, including ATF4, ATF5, EIF2S2, SARS1, PHGDH, PSAT1, ASNS and PYCR1, together with suppression of genes involved in fatty acid oxidation and mitochondrial metabolism. Hierarchical clustering and gene set enrichment analysis further showed activation of mTORC1 signaling and amino acid biosynthesis, with suppression of fatty acid metabolism (Extended Data Fig. 5a-c). Principal component analysis revealed clear separation by genotype, with sex contributing predominantly to the second principal component (Extended Data Fig. 5d). Integration of transcriptomic and metabolomic datasets identified glycine, serine and threonine metabolism as the most prominently altered pathway (Extended Data Fig. 5e).

To determine whether these transcriptional changes were accompanied by altered metabolic flux, we performed ^13^C-glucose metabolomic tracing (Fig. 4e) in WT and cPHB1KO hearts at 6 weeks following Cre induction. cPHB1KO hearts had dramatically increased levels of M+3 serine and M+2 glycine enrichment compared with WT (Fig. 4f), indicating increased glucose-derived carbon flux into *de novo* serine and glycine synthetic pathway. Total glycolytic flux was not significantly different between WT and cPHB1KO, as ^13^C enrichment of pyruvate, lactate, α-ketoglutarate, glutamine and asparagine was unchanged (Extended Data Fig. 6). Consistent with the ^13^C tracer analysis, metabolite profiling showed reduced glutamine abundance in cPHB1KO hearts of both sexes, with increased abundance of serine, glycine, asparagine and leucine (Fig. 4g). Together, these findings suggest PHB complex ablation activates mTORC1 and ISR signaling to drive pathogenesis of cardiomyopathy via altered glucose and amino acid metabolism in the cPHB1KO hearts.

### Cardiomyocyte PHB complex ablation causes impaired cardiac autophagy that is not responsive to fasting

Given the well-established role of mTORC1 in regulating autophagy initiation and that derangements in autophagy are known to promote cardiomyopathy ^28–31^, we examined whether sustained mTORC1 signaling altered autophagy in cPHB1KO hearts. Immunoblot analysis of left ventricular extracts prepared from mice euthanized in the fed state showed that phosphorylation of ULK1 at Ser757 was unchanged at 3 and 6 weeks but increased significantly at 9 and 12 weeks following Cre induction with tamoxifen. The LC3-II/LC3-I ratio was preserved through 9 weeks and reduced at 12 weeks, whereas p62 accumulated progressively from 6 to 12 weeks (Fig. 5a, b). TEM further showed accumulation of autolysosomes and lipid droplets in cPHB1KO hearts at 6 weeks (Fig. 5c).

**Fig. 5:**
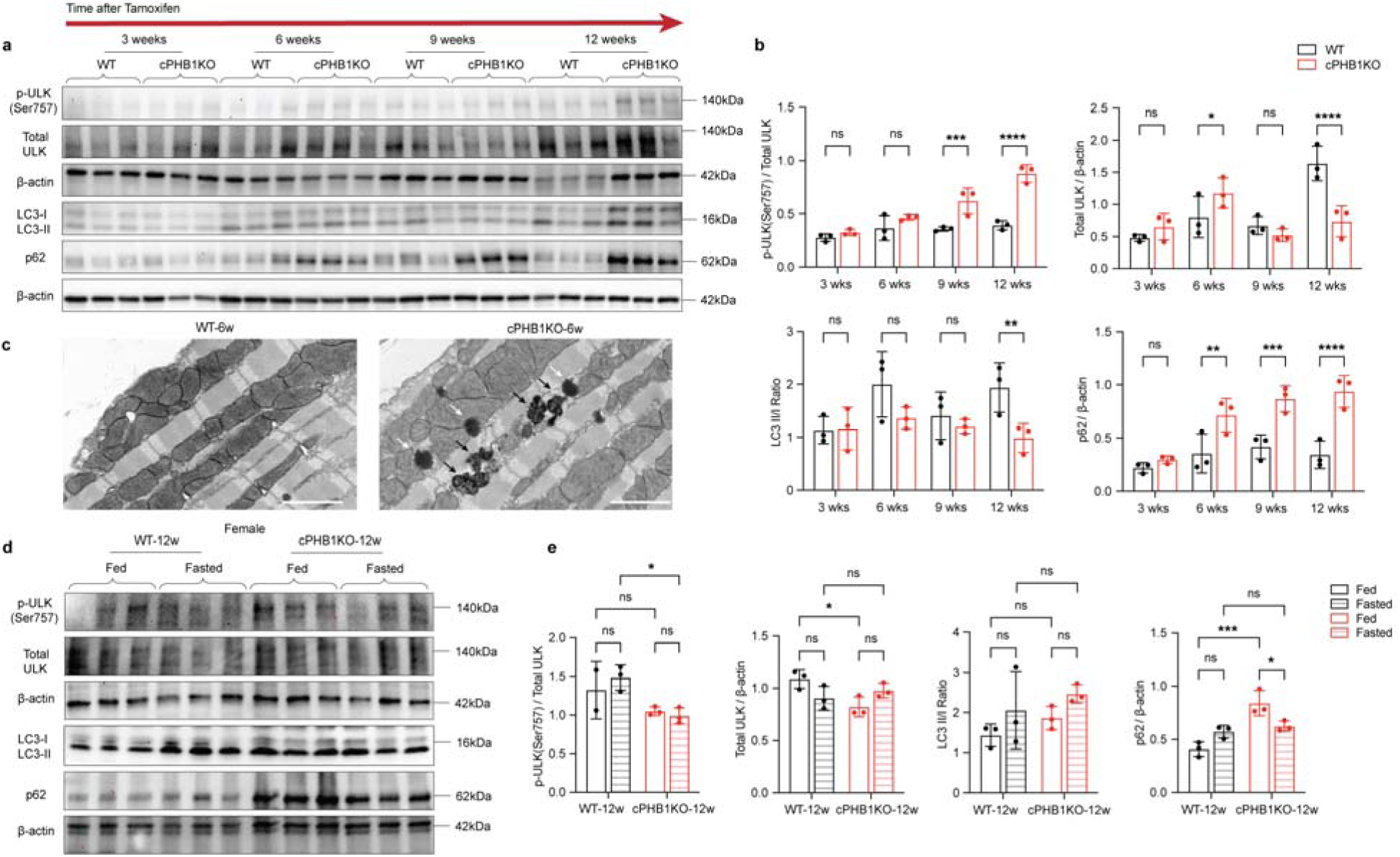
Impaired autophagy after loss of PHB complex in the heart. **a**, Western blot of autophagy markers, including phospho-ULK1 (Ser757) (p-ULK Ser757), total ULK1, LC3-I and LC3-II, and p62, in left ventricular tissue extracts from female WT and cPHB1KO mice at 3, 6, 9 and 12 weeks post-tamoxifen. Gels/blots were processed in parallel. Data are representative of three biological replicates. β-actin was used as the loading control. b, Quantification of the immunoblots in a. Data are mean ± s.d. Statistical significance was determined by two-way ANOVA with Šidák’s multiple-comparisons test. c, Representative transmission electron microscopy (TEM) images of left ventricle from WT and cPHB1KO mice at 6 weeks post-tamoxifen (WT-6w and cPHB1KO-6w). White arrowheads indicate lipid droplets and black arrowheads indicate autolysosomes. Magnification, ×20,000. Scale bars, 2 µm. d, Western blot of the same autophagy markers in left ventricle from female WT and cPHB1KO mice at 12 weeks post-tamoxifen (WT-12w and cPHB1KO-12w) under fed and fasted conditions. Gels/blots were processed in parallel. Data are representative of three biological replicates. β-actin was used as the loading control. e, Quantification of the immunoblots in d. Data are mean ± s.d. Statistical significance was determined by two-way ANOVA with Tukey’s multiple-comparisons test. For all panels, \**P* < 0.05, \*\**P* < 0.01, \*\*\**P* < 0.001, \*\*\*\**P* < 0.0001. n.s., not significant.

To determine whether hyperactivated mTORC1 retained its sensitivity to nutrient deprivation in the cPHB1KO hearts, we subjected a subset of mice to an 18-hour fast at 12 weeks following Cre induction. Fasting suppressed mTORC1 signaling in WT hearts but produced only a limited response in cPHB1KO hearts. Phosphorylated mTOR and 4EBP1 remained elevated relative to WT under fasting conditions, although p-4EBP1 decreased within the cPHB1KO group (Extended Data Fig. 7a–d). Total mTOR and 4EBP1 abundance were largely unchanged. Similarly, p62 remained elevated in cPHB1KO hearts under both fed and fasted conditions in females (Fig. 5d,e) and males (Extended Data Fig. 7e,f), while the LC3-II-to-LC3-I ratio showed no response to fasting in cPHB1KO. Male cPHB1KO hearts also retained elevated p-ULK1 Ser757 under both fed and fasted states (Extended Data Fig. 7f). These findings indicate impaired autophagic homeostasis and a significantly blunted response to fasting in cPHB1KO hearts.

### mTORC1 inhibition blunts pathogenesis of cardiomyopathy in female cPHB1KO mice

We next tested whether mTORC1 inhibition with chronic rapamycin treatment could attenuate progression of cardiomyopathy in cPHB1KO mice. After 9 weeks of rapamycin treatment (2 mg/kg i.p., q.o.d starting 1 week after Cre induction with tamoxifen, Fig. 6a), cPHB1KO mice had reduced levels of total and phosphorylated mTOR in their hearts, and sharply reduced total and phosphorylated RPS6, while p-4EBP1 (Ser64) was unchanged (Fig. 6b–d). Interestingly, rapamycin significantly reduced heart weight-to-tibia length ratio only in female cPHB1KO mice, with no corresponding effect in males (Fig. 6e,f). Serial echocardiography showed that rapamycin preserved ejection fraction in female cPHB1KO mice, whereas male cPHB1KO mice showed no functional improvement (Fig. 6g). Rapamycin did not significantly alter p-ULK1 Ser757, total ULK1, LC3-II/LC3-I or p62 in female cPHB1KO hearts (Extended Data Fig. 8), indicating autophagy was unresponsive to mTORC1 inhibition.

**Fig. 6:**
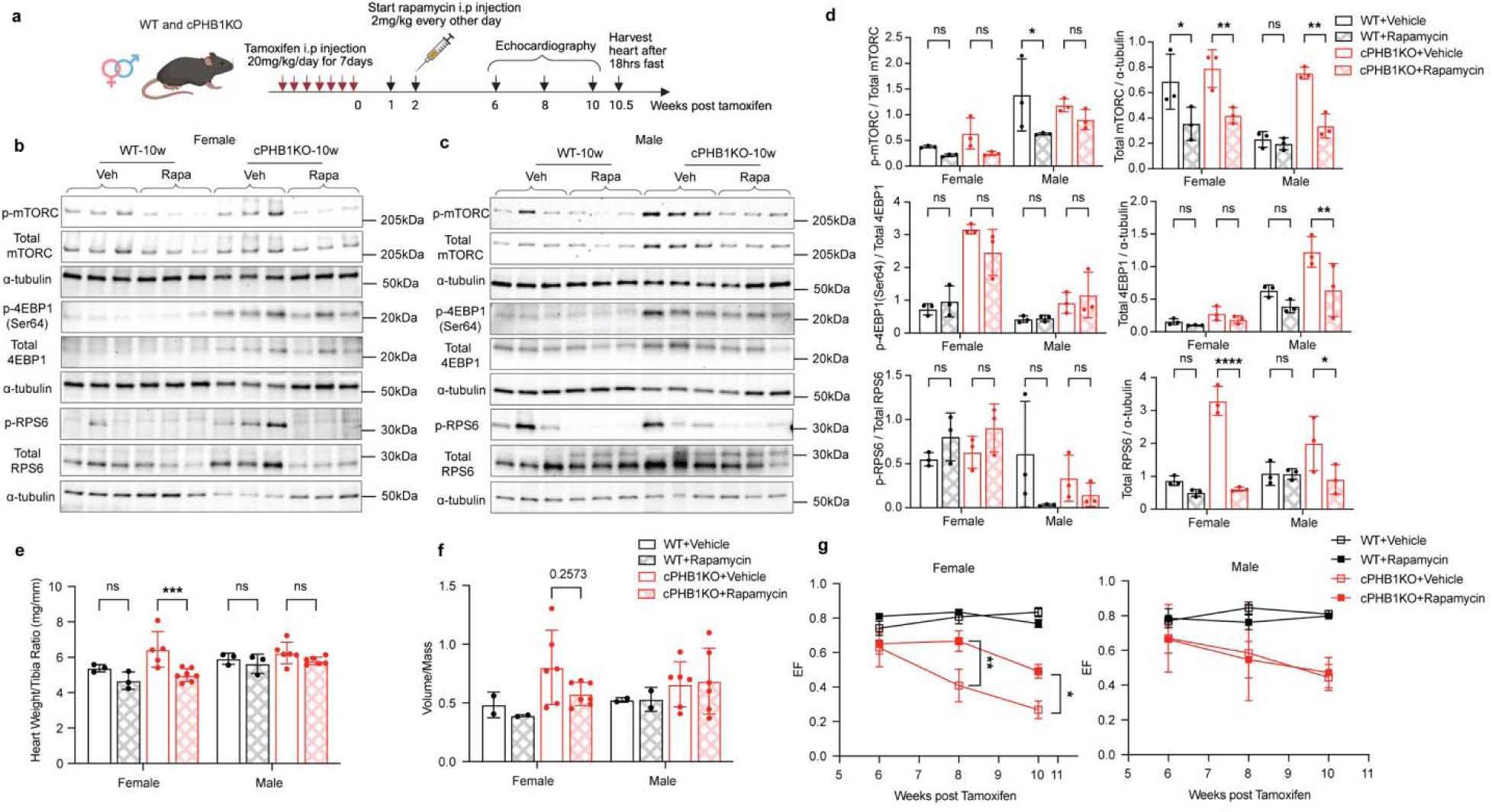
mTORC1 inhibition by rapamycin mitigates progression of dilated cardiomyopathy in sex-dimorphic manner. **a**, Schematic of rapamycin treatment and experimental time points in WT and cPHB1KO mice. b**,c**, Western blot of mTOR-pathway proteins phospho-mTOR (p-mTOR), total mTOR, phospho-4EBP1 (Ser64) (p-4EBP1 Ser64), total 4EBP1, phospho-RPS6 (p-RPS6) and total RPS6 in left ventricular tissue extracts from female (b) and male (c) WT and cPHB1KO mice at 10 weeks post-tamoxifen (WT-10w and cPHB1KO-10w), treated with vehicle (Veh) or rapamycin (Rapa). Mice were fasted 18 hours prior to euthanasia. Gels/blots were processed in parallel. Data are representative of three biological replicates. α-tubulin was used as the loading control. d, Quantification of the immunoblots in b and c. Data are presented as mean ± s.d. Significance was determined by two-way ANOVA with Tukey’s multiple-comparisons test. e, Heart weight to tibia length ratio in female and male WT and cPHB1KO mice treated with vehicle or rapamycin. Data are mean ± s.d. Female: WT+Vehicle, *n =* 3; WT+Rapamycin, *n =* 3; cPHB1KO+Vehicle, *n* = 5; cPHB1KO+Rapamycin, *n* = 7. Male: WT+Vehicle, *n* = 3; WT+Rapamycin, *n* = 3; cPHB1KO+Vehicle, *n* = 6; cPHB1KO+Rapamycin, *n* = 6 mice. Significance was determined by two-way ANOVA with Tukey’s multiple-comparisons test. f, Left ventricular volume-to-mass ratio in the same groups. Data are mean ± s.d. Female: WT+Vehicle, *n* = 2; WT+Rapamycin, *n* = 2; cPHB1KO+Vehicle, *n* = 6; cPHB1KO+Rapamycin, *n* = 7. Male: WT+Vehicle, *n* = 2; WT+Rapamycin, *n* = 2; cPHB1KO+Vehicle, *n* = 6; cPHB1KO+Rapamycin, *n* = 6 mice. Significance was determined by two-way ANOVA with Tukey’s multiple-comparisons test. g, Left ventricular ejection fraction (EF) measured by echocardiography at 6, 8 and 10 weeks post-tamoxifen in female (left) and male (right) mice. Data are mean ± s.e.m. Female: WT+Vehicle, *n* = 2; WT+Rapamycin, *n* = 3; cPHB1KO+Vehicle, *n* = 5; cPHB1KO+Rapamycin, *n* = 7. Male: WT+Vehicle, *n* = 2; WT+Rapamycin, *n* = 2; cPHB1KO+Vehicle, *n* = 6; cPHB1KO+Rapamycin, *n* = 6 mice. Significance was determined by unpaired two-tailed Welch’s t-test at 8 and 10 weeks. For all panels, \**P* < 0.05, \*\**P* < 0.01, \*\*\**P* < 0.001, \*\*\*\**P* < 0.0001. n.s., not significant.

## Discussion

Molecular pathways by which cardiac metabolism contributes to maladaptive structural and functional remodeling has remained a vexing topic of HF research for many decades. On the one hand, the failing heart exhibits many hallmarks of energy deprivation and metabolic ‘exhaustion,’ as elegantly described nearly 20 years ago by Neubauer ^32^. While evidence of this phenotype can undeniably be found in biopsies of HF patients and experimental models (e.g., loss of mitochondria, decreased ATP, etc), the reality is that HF most often manifests itself in patients that are obese and often diabetic, where the myocardium is in a state of nutrient overload rather than deprivation. Even more perplexing is that the heart is the most metabolically ‘flexible’ of all organs in the body and can readily use any source of carbon presented to it for fuel, at any time. Recent theories by Taegtmeyer others proffers a more nuanced explanation for this paradox by suggesting that the problem is likely to be driven by a ‘reprogramming’ of metabolism in the heart that occurs either in concert with, or shortly after, hemodynamic stress or injury ^2, 5, 33^. Thus, it is the metabolic adaptations that follow the stress, and the molecular mechanisms driving these adaptations, that are ultimately responsible for HF pathogenesis. These mechanisms remain unresolved to this day.

Here, we have presented an intriguing model of metabolic cardiomyopathy that we believe may provide some insights on these mechanisms. At the nexus of this cardiomyopathy lie the mitochondrial PHB complex and mTORC1, both of which are well established mediators of growth factor and nutrient signaling, cell proliferation and glucose metabolism, all of which are key drivers of cancer pathogenesis ^34, 35^. Specifically, our study illustrates that, when accompanied by derangements in cardiomyocyte ROS and Ca^2+^ handling, reprogramming of glucose metabolism into anaplerotic and biosynthetic pathways leads to severely dilated myocardium and rapid mortality.

We find that cardiomyocyte-specific PHB1 deletion induced the loss of the PHB1,2 complex, upregulated the integrated stress response (ISR) and unrestrained mTORC1 signaling, and increased glucose-derived carbon incorporation into biosynthetic pathways, particularly *de novo* serine and glycine biosynthesis (Fig. 7). These early changes preceded progressive mitochondrial respiratory failure, impaired autophagic homeostasis and lethal dilated cardiomyopathy (DCM). Notably, metabolic reprogramming occurred while mitochondrial respiration and ATP production remained largely preserved (Fig. 2), indicating that altered glucose-derived carbon allocation is an early response to PHB loss rather than simply a consequence of bioenergetic failure. Thus, despite their limited proliferative capacity, cardiomyocytes are capable of engaging an anabolic program resembling the Warburg-like phenotype observed in proliferating cells ^36^.

The most distinctive metabolic feature of pre-symptomatic cPHB1KO hearts was increased ^13^C-glucose incorporation into serine and glycine (i.e. purine synthesis) pathway, accompanied by induction of ATF4, PHGDH, PSAT1 and other genes involved in amino acid biosynthesis. Integrated transcriptomic and metabolomic analyses similarly identified glycine, serine and threonine metabolism as a major pathway altered by PHB1 deficiency. Serine synthesis is increasingly recognized as an important component of cardiac remodeling, although its functional consequences appear context dependent. YAP-dependent activation of serine biosynthesis has been linked to cardiomyocyte hypertrophy ^37^, whereas increased PHGDH activity and serine one-carbon metabolism have also been reported to protect against cardiac dysfunction ^38, 39^. A recent report indicated enhanced cardiac hypertrophy and dilated cardiomyopathy in PHGDH^+/-^ mice following transverse aortic constriction (TAC) ^40^. In cPHB1KO hearts, increased *de novo* serine/glycine flux occurred before detectable contractile dysfunction and in the setting of sustained ROS production (Fig. 3). Whether serine/glycine biosynthesis contributes directly to disease progression or represents an early compensatory mechanism therefore remains unresolved and will require further investigation. In any case, the reduction in myocardial glutamine and changes in asparagine and leucine further support broad remodeling of amino acid metabolism early in cPHB1KO hearts, whereas suppression of fatty acid oxidation genes suggests early transcriptional remodeling of oxidative substrate metabolism.

These metabolic changes in cPHB1KO hearts were accompanied by sustained alterations in mTOR signaling. We recently showed that hepatocyte PHB1 deficiency (i.e., PHB1^+/-^) increases mTORC1 signaling and promotes de novo amino acid and lipid biosynthesis in the liver ^18^, suggesting that regulation of anabolic signaling by the PHB complex is similar across metabolically active tissues. In the heart, PHB1 loss produced sustained phosphorylation of 4EBP1 at Ser64 together with transcriptional enrichment of mTORC1 regulated genes. However, the lack of parallel activation of p70S6K (Fig. 4) and the limited effect of rapamycin on 4EBP1 Ser64 phosphorylation (Fig. 6) suggest redundancies must be responsible for hypertrophy in these hearts. A recent report indicates that cardiomyocyte 4EBP1 phosphorylation can be directly controlled by both mTORC1 and ERK ^23^, raising the possibility that PHB deficiency engages multiple pathways controlling protein translation and synthesis.

Also consistent with hyperactive mTORC1 signaling, cPHB1KO hearts progressively accumulated p62, developed increased inhibitory ULK1 phosphorylation and eventually showed reduced LC3 lipidation. These abnormalities were accompanied by accumulation of autolysosomes and lipid droplets and were only minimally responsive to fasting. It is noteworthy that a recent study showed that cardiomyocyte-specific PHB2 knockdown leads to impairments in fatty acid oxidation and lipid droplet accumulation ^41^. Together, these findings support progressive disruption of autophagic homeostasis, although direct autophagic flux measurements will be required to distinguish altered autophagosome formation from impaired lysosomal clearance.

Changes in mitochondrial Ca²⁺ handling also were observed in cPHB1KO hearts prior to remodeling (Fig. 3). Specifically, at 6 weeks post Cre induction, while cPHB1KO hearts were structurally and functionally normal, cardiac mitochondria in these mice had substantially higher rates of Ca²⁺ uptake in both sexes and increased Ca²⁺ retention capacity in females despite largely preserved respiration, ATP production and respiratory complex abundance. An interesting finding with these experiments was that acute supplementation of the mitochondria with recombinant PHB1, alone or with PHB2, attenuated these abnormalities. This suggests PHBs may physically interact with transporters responsible for regulating mitochondrial Ca²⁺ uptake (e.g., MICU1, MCU). Immunoblot analysis indicated that MICU1 and MCU abundance remained largely preserved with PHB complex ablation, but declined later as cardiac dysfunction progressed, suggesting that remodeling of the MCU complex is more likely to amplify rather than initiate the early Ca²⁺ handling phenotype. Increased cellular ROS and early mitochondrial structural abnormalities further indicate that mitochondrial stress develops first, before measurable respiratory failure ensues. These observations place altered Ca²⁺ handling and redox homeostasis among the earliest mitochondrial consequences of PHB complex loss and raise the possibility that these are early stressors that are compounded by metabolic reprogramming to ultimately drive structural remodeling in the heart.

A notable feature of the cPHB1KO phenotype was its sexual dimorphism. Female cPHB1KO mice progressed more rapidly to mortality and exhibited greater abnormalities in mitochondrial Ca²⁺ handling and ROS, yet also showed greater benefit from rapamycin. Rapamycin attenuated the decline in ejection fraction and reduced cardiac remodeling in female cPHB1KO mice, whereas little benefit was observed in males (Fig. 6). The functional improvement in females was accompanied by a pronounced reduction in RPS6 abundance but not by normalization of 4EBP1 phosphorylation or the measured autophagy markers. Thus, the sex-specific benefit cannot be attributed to restoration of autophagic homeostasis and may instead reflect differential dependence on mTORC1-mediated protein translation/synthesis between sexes. Indeed, sex-specific differences in mTOR signaling and cardiac remodeling have been described previously ^42, 43^, but the mechanisms underlying the response observed here remain unresolved. Interestingly, a similar sex-dimorphic effect of hepatocyte PHB deficiency was found in our recent study. In hepatocytes, PHB1 deficiency led to hyperactive mTORC1 signaling, but it was only males that were affected and showed evidence of fatty liver and insulin resistance, while females were phenotypically normal. Inhibition of mTORC1 with Torin1 reversed the metabolic abnormalities in the male mice ^18^. A sex dimorphic role for PHBs in adipose tissue expansion and fatty liver disease has been reported by others ^44^, further supporting a strong sex-dependence for the biological effects of PHBs. Taken together, the sex dimorphic regulation of PHBs and mTORC1 signaling remains an area that clearly demands further investigation.

Collectively, our findings position the PHB complex as a key regulator of cardiac metabolic homeostasis and reveal novel insights on mTORC1 regulation of anabolic signaling and serine/glycine metabolism in pathogenesis of cardiomyopathy. Given that both PHBs ^13, 14^ and mTORC1 ^45, 46^ remain compelling drug targets for a wide variety of disorders associated with aging and obesity, we anticipate this study will inform further investigations of these targets and lead to new therapies for cardiomyopathy and other forms of heart disease in near future.

## Acknowledgements

Authors wish to extend their gratitude to Dr. William Paradee and the staff at the University of Iowa Viral Vector core facility for their assistance generating *Phb1^fl/fl^* mice, and all scientific staff at the University of Iowa Genomics, Metabolomics, Central Microscopy, and Metabolic Phenotyping core facilities for their resources and expertise.

## Funding

This work was supported by funding from American Heart Association Career Development Award #935054 (Q.S), American Heart Association Strategically Focused Research Network award 20SFRN35200003 (E.J.A.), Department of Veteran Affairs I01BX006139 (L.S.), and National Heart Lung Blood Institute awards HL176940, HL177305, HL181428 (L.S.) and HL167087 (E.J.A.).

## Conflict of Interest

The authors have no conflicts of interest to declare.

## Methods

### Animals

Inducible cardiomyocyte-specific PHB1 knockout (cPHB1KO) mice were generated by crossing mice carrying floxed PHB1 alleles with transgenic mice expressing tamoxifen-inducible MerCreMer recombinase under the control of the α-myosin heavy-chain promoter (αMHC-MerCreMer; MCM ^24^). The MCM mice, maintained on a C57BL/6 background, were provided by Long-Sheng Song. Subsequent breeding generated PHB^fl/fl^, MCM^+/-^ mice, referred to throughout the manuscript as cPHB1KO. PHB^-/-^, MCM^+/-^ mice were used as controls to account for potential cardiotoxic effects of tamoxifen-induced Cre recombinase as described previously ^48^. Male and female mice aged 12–16 weeks were randomly assigned to experimental groups. Tamoxifen was dissolved in corn oil and administered by intraperitoneal injection at 20 mg/kg/day for seven consecutive days. Rapamycin (MedChemExpress, HY-10219) was dissolved in a filter-sterilized solution containing 1.2% DMSO, 5% PEG-400, 5% Tween-80 and sterile saline and administered by intraperitoneal injection at 2 mg/kg every other day, beginning 2 weeks after tamoxifen. Mice were maintained on standard chow unless otherwise indicated. All animal procedures were approved by the Institutional Animal Care and Use Committee of the University of Iowa and were conducted in accordance with the National Institutes of Health Guide for the Care and Use of Laboratory Animals.

### Echocardiography

Echocardiographic evaluations and analyses were performed in the University of Iowa Cardiology Animal Phenotyping Core Laboratory. Hair was removed from the left side of the chest before imaging. Conscious mice were restrained by the nape and positioned in the left lateral recumbent position. Two-dimensional and M-mode images of the left ventricle were acquired in the short- and long-axis views using a Vevo 2100 imaging system equipped with a 30-MHz linear-array transducer (Vevo 2100; Visual Sonics, Toronto, ON, Canada). Image acquisition and analysis were performed by an experienced echocardiographer who was blinded to mouse genotype.

### Isolation of cardiac mitochondria

Hearts were removed from mice under deep anesthesia and quickly rinsed in ice-cold phosphate-buffered saline. Atria were removed, and the ventricles were blotted dry, minced and processed within approximately 2 min of excision. Minced ventricular tissue was incubated on ice for 2 min in mitochondrial isolation medium (MIM; 300 mM sucrose, 10 mM Na-HEPES and 0.2 mM EDTA, pH 7.2) containing trypsin (Sigma-Aldrich, T0303). Digestion was terminated by adding trypsin inhibitor (Sigma-Aldrich, T9003). Tissue was resuspended in 5 ml MIM containing 0.1% fatty-acid-free bovine serum albumin (BSA) and homogenized with ten manual strokes using a polytetrafluoroethylene pestle. The homogenate was centrifuged at 600 × g for 10 min at 4 °C. The supernatant was collected and centrifuged at 8,000 × g for 15 min at 4 °C. The mitochondrial pellet was resuspended in MIM containing 0.1% fatty-acid-free BSA and washed by a second centrifugation at 8,000 × g for 15 min at 4 °C. EGTA was added to all digestion and homogenization buffers at a final concentration of 1 mM. The final mitochondrial pellet was resuspended in 40–50 µl MIM without EGTA. For protein quantification, 2 µl mitochondrial suspension was diluted in 98 µl water, and protein concentration was measured using the Pierce BCA Protein Assay Kit (Thermo Fisher Scientific). Mitochondria used for functional assays were prepared on the day of analysis. Mitochondria used for immunoblotting were stored at −80 °C.

### Preparation of permeabilized cardiac myofibers

The procedure for preparing permeabilized cardiac myofibers from dissect myocardium has been detailed previously by our group ^49–52^. In brief, mice were anesthetized with ketamine/xylazine and hearts were rapidly excised. A section of left ventricular myocardium was placed immediately in ice-cold Buffer X (7.23 mM K_2_EGTA, 2.77 mM CaK_2_EGTA, 0.5mM DTT, 20 mM Imidazole, 20 mM Taurine, 5.7 mM ATP, 14.3 mM phosphocreatine, 6.56 mM MgCl_2_-6H_2_O, and 50 mM MES, pH 7.1). Small myofiber bundles weighing approximately 1–2 mg was separated under a dissecting microscope using fine forceps. Bundles were permeabilized in Buffer X containing 50 µg/mL saponin for 40 min. The permeabilized myofibers were subsequently transferred to ice-cold Buffer Z-lite (105 mM K-MES, 30 mM KCl, 10 mM KH_2_PO_4_, 5 mM MgCl_2_-6H_2_O, 0.5 mg/mL BSA). Buffer Z-lite was supplemented with 20 µM blebbistatin to prevent Ca^2+^-independent contraction. Bundles were maintained on a rocker at 4 °C until analysis.

### Mitochondrial *J*O_2_, *J*ATP, *J*H_2_O_2_ and Ca^2+^ uptake measurements

All experiments were conducted at 30 °C in Buffer Z-lite. Oxygen consumption rates (*J*O_2_) were measured using the Oroboros O_2_K Oxygraph system. The assay buffer for myofibers is Buffer Z-lite, supplemented with 20 mM creatine monohydrate and 20 µM (−)-Blebbistatin (Sigma, B0560), while Buffer Z-lite with 1mM EGTA is used for isolated mitochondria. Oxidative substrates (5 mM Pyruvate, 50 µM Palmitoylcarnitine, 5 mM Glutamate, 2 mM Malate, 5 mM Succinate, 500 µM ADP) were added in a specific order. Mitochondrial ATP production (*J*ATP) was assessed by coupling ATP hydrolysis to NADPH production and subsequent autofluorescence, using a custom tandem oxi-fluorometer approach developed and validated in our laboratory ^50^. To ensure coupling of ATP hydrolysis to NADPH release, the following reagents were added to the assay media: 4 mM D-glucose, 1.7 U/mL glucose-6-phosphate dehydrogenase and 3.4 U/mL hexokinase (G6PDH/HK, Roche), 200 µM nicotinamide adenine dinucleotide phosphate (NADP+, Sigma, N0505) and 50 µM P1,P5-Di (Adenosine-5′) pentaphosphate (Ap5A) (Sigma, D4022) to inhibit adenylate kinase and confirm that the measured ATP production was exclusively mitochondrial.

Mitochondrial H_2_O_2_ production (*J*H_2_O_2_) and Ca^2+^ uptake studies were conducted using a spectrofluorometer (Photon Technology Instruments, Birmingham, NJ) equipped with a thermal-jacketed cuvette chamber. All experiments were carried out at 37 °C. For mitochondrial H_2_O_2_ measurements, 1 mL Buffer Z-lite was used, containing 10 μM Amplex Red, 1 U/mL horseradish peroxidase (HRP), 5 U/mL superoxide dismutase (SOD), 5 mM pyruvate, and 2 mM malate. The cuvette was placed in the fluorometer to establish a baseline for approximately 5 minutes. Subsequently, 5 mM succinate was added to the cuvette to initiate reverse electron transport flow and induce ROS production. For the measurement of Ca^2+^ uptake, 100-200 μg isolated mitochondria was added into 1 mL Buffer Y (250 mM sucrose, 10 mM Tris-Cl, 20 mM Tris base, 5 mM KH_2_PO_4,_ 0.5 mg/mL BSA), containing 5 μM Calcium Green 5-N, 5 mM pyruvate, 2 mM malate, and 5 mM succinate. Additionally, 0.5 µM EGTA was included to bind any remaining Ca^2+^. Sequential injections of 10 μL of 1 mM Ca^2+^ (CaCl_2_) were administered, and the absorption of Ca^2+^ was monitored up to the point where the mitochondrial permeability transition pore was activated. A 40 μL of 10 mM bolus of CaCl_2_ was added to saturate the probe at the experiment’s end. The changes in free calcium concentration were determined using the established K_d_ for Calcium Green 5-N along with the minimum (F_min_) and maximum (F_max_) fluorescence recorded in each experiment, applying the formulas developed by Tsien ^53^.

### Mitochondrial membrane micro-fluidity

Mitochondrial membrane micro-fluidity was assessed using merocyanine 540 (MC540). Freshly isolated mitochondria (100 µg protein) were suspended in 1 ml MIM without BSA in a 3.5-ml quartz cuvette with a 1 cm path length. MC540 was added to a final concentration of 100 nM from a 100 µM stock prepared in DMSO. Fluorescence measurements were performed using a spectrofluorometer (Photon Technology Instruments, Birmingham, NJ) equipped with a thermal-jacketed cuvette chamber at 30 °C. Samples were equilibrated for 2–3 min before measurement. Emission spectral scans were recorded from 540 to 660 nm using an excitation wavelength of 495 nm. MC540 fluorescence intensity at the emission maximum of approximately 580 nm was used to compare mitochondrial membrane fluidity between experimental groups.

### Isolation of adult primary cardiomyocytes from mice

Mice were anesthetized with ketamine/xylazine and hearts were rapidly excised and placed in ice-cold Ca^2+^-free Tyrode’s buffer. Hearts were cannulated through the aorta and retrogradely perfused using a Langendorff apparatus with digestion buffer containing collagenase and protease. After about 20 minutes, softened hearts were minced, gently triturated, and filtered through nylon to isolate cardiomyocytes. The cells were allowed to settle, the supernatant was removed, and the cardiomyocytes were resuspended in calcium-containing Tyrode’s buffer. After 15 minutes, the cardiomyocytes sank into the bottom of the tube and were collected for further analysis.

### Transmission electron microscopy

Hearts were dissected from anesthetized mice cannulated through the aorta, and perfused on a Langendorff apparatus with saline for 10 minutes to flush and eliminate residual blood. The perfusion was then switched to a 2% EM-grade glutaraldehyde solution for fixation, lasting between 20 and 30 minutes. During this process, a beaker was used to recirculate the fixation solution while simultaneously submerging the heart in it, ensuring both internal and external fixation. Following fixation, a 1 cm² piece of myocardium was excised from the left ventricle and further sectioned into 1 mm² pieces. These tissue samples were kept in the fixation solution overnight at 4 °C and subsequently post-fixed in 1% osmium tetroxide for 2 hours at room temperature. The osmicated samples were then dehydrated in a series of graded acetone solutions and embedded in epoxy resin. At the University of Iowa Central Microscopy Core Facility, ultra-thin sections (600–800 Å) were cut from the resin blocks. Finally, the grids were stained with lead citrate and imaged using a Hitachi transmission electron microscope equipped for digital image acquisition.

### *In situ* confocal imaging of whole myocardium

All procedures have been described previously by Dr. Song’s group ^54, 55^. In brief, hearts were removed from anesthetized mice and perfused via a retrograde Langendorff system with Krebs-Henseleit (KH) solution (120 mM NaCl, 24 mM NaHCO_3_, 11.1 mM glucose, 5.4 mM KCl, 1.8 mM CaCl_2_, 1 mM MgCl_2_, 0.42 mM NaH_2_PO_4_, 10 mM taurine, 5 mM creatine) for 10 minutes. Subsequently, the perfusion was switched to KH solution containing 2 μM tetramethylrhodamine ethyl ester (TMRE, Invitrogen T669) and 10 μM CM-H2DCFDA (DCF, General Oxidative Stress Indicator, Invitrogen C6827) at room temperature for 20 minutes. TMRE was used to measure mitochondrial membrane potential, while DCF was used for reactive oxygen species (ROS) detection. After loading TMRE and DCF, hearts were transferred to a confocal microscope system connected to another Langendorff apparatus set to 37 °C with a perfusion of KH solution oxygenated with 95% O2 and 5% CO2. The heart was positioned and set to sinus rhythm in a recording chamber for in situ confocal imaging of the TMRE and DCF fluorescence in the epicardial myocytes. To minimize motion artifacts during imaging, 10 μM (±)-Blebbistatin (Cayman, No.13186) was added to the perfusion solution. Imaging of TMRE and DCF was performed using tandem excitation at 488 and 561 nm, with emission collected at 500–545 and >560 nm, respectively. The heart was allowed to stabilize before acquiring control images. Confocal imaging was conducted with a confocal microscope (LSM510, Carl Zeiss MicroImaging Inc.) equipped with a 63× (N.A. = 1.4) oil immersion lens. Ten images from different regions of the left ventricular wall were captured, and intensity values from each ventricle were analyzed using ImageJ software and averaged to represent the overall TMRE and DCF fluorescence intensity for each ventricle.

### Blue native PAGE, Coomassie staining and blotting

The Blue Native PAGE method was adapted from previous reports ^56, 57^. Mitochondrial pellets were resuspended in ACA buffer (0.75 M 6-aminocaproic acid, 50 mM bis-tris, pH 7.0) containing freshly added protease inhibitors. A small aliquot of the suspension was further diluted and lysed in 1:50 ratio in water, and the protein concentration was determined using a BCA assay. To normalize the concentration, samples were adjusted to 2.5 mg/mL with ACA buffer. Digitonin (high purity, Millipore) was added at a 4:1 weight ratio to mitochondrial protein, and the mixture was incubated on ice for 20 minutes with gentle vortexing every 5 minutes. Following centrifugation at 20,000 × g for 30 minutes, the supernatant was mixed with Coomassie Brilliant Blue G-250 (CBBG, Sigma B0770, 1:4 g:g, digitonin: CBBG) and glycerol (Final 5% volume), then loaded onto a native 3-12% or 4-16% gel (Invitrogen). High molecular weight markers (GE Healthcare) were prepared in 100 uL ACA plus protease inhibitors and 5 uL of 5% CBBG and loaded at the first and last lanes. The gel was run at 150V in a cold room using a dark blue cathode buffer with 0.02% CBBG until the dye front reached halfway; then switched to a light blue cathode buffer with 0.0005% CBBG and increased to 250V for an additional two hours. For Coomassie staining, gels were stained in staining buffer (0.5% CBBG in 50% methanol, 10% acetic acid) on a rocker for 5 minutes, destained in 40% methanol / 10% acetic acid for 4 times every 15 minutes and destaining continued overnight if needed with Kim wipes to absorb excess stain until the bands are very clean. For blotting, proteins were transferred to a PVDF membrane using an Invitrogen chamber or Bio-Rad system in NuPAGE transfer buffer at 20V for 2 to 5 hours without methanol, ethanol, or SDS. Membranes were fixed in 8% acetic acid, rinsed in water, air-dried, re-wetted in methanol, and blocked with 5% BSA in TBS-0.05% tween (TBST). Primary and secondary antibodies were applied, and the blot was developed using enhanced chemiluminescence and captured on an iBright Imaging system (ThermoFisher Scientific, Inc., Waltham, MA).

### Immunoblot analysis

Heart tissue left ventricles (50mg) were homogenized in 1mL of ice-cold tissue lysis buffer (50 mM HEPES, 150 mM NaCl, 1% Triton X-100, 2 mM EGTA, 1 mM MgCl_2_), supplemented with 0.5 mM DTT and 1% Protease/phosphatase inhibitor. Meanwhile, isolated adult cardiomyocytes were lysed in 0.5-1 mL RIPA buffer (Thermo Fisher Scientific, Waltham, MA) containing the Complete protease inhibitor cocktail (Roche). The tissue or cell lysates were centrifuged at 20,000 × g for 15 minutes at 4 °C. Protein concentration was determined using the BCA system, and protein lysates were denatured by adding LDS buffer and heating for 10-20 minutes at 65 °C. Protein samples (10 μg/lane) were separated on a 4–12% BisTris Precast gel (Bio-Rad) and then transferred to a PVDF membrane (Bio-Rad). The membranes were incubated with primary antibodies overnight at 4 °C as follows: PHB1 (Abcam, ab75766, 1:10,000), PHB2 (Cell Signaling Technology, E1Z5A, 1:2,000), total OXPHOS rodent antibody cocktail (Abcam, ab110413, 1:3,500), MCU (Cell Signaling Technology, 14997, 1:1,000), MICU1 (Sigma-Aldrich, HPA037480, 1:1,000), β-actin (Abcam, ab8227, 1:4,000), phospho-4EBP1 (Cell Signaling Technology, 9451, 1:1,000), total 4EBP1 (Invitrogen, AH01382, 1:500), phospho-p70S6K (Invitrogen, MA5-15202, 1:1,000), total p70S6K (Proteintech, 14485-1-AP, 1:1,000), phospho-ULK1 (Ser757) (Cell Signaling Technology, 6888, 1:1,000), total ULK1 (Cell Signaling Technology, 8054, 1:1,000), LC3A/B (Novus Biologicals, NB600-1384, 1:3,000), p62 (Sigma-Aldrich, P0067, 1:1,000), phospho-mTOR (Ser2448) (Cell Signaling, 5536S, 1:1000), total mTOR (Cell Signaling, 2983S, 1:1000), phospho-RPS6 (Ser240/244) (Cell Signaling, 5364T, 1:1000), total RPS6 (Cell Signaling, 2317S, 1:1000) and α-tubulin (Abcam, AB52866, 1:1000).Secondary antibodies were HRP-conjugated goat anti-mouse-IgG (Jackson Immunoresearch, cat #115-035-003, 1:2000) and HRP-conjugated anti-rabbit-IgG (Jackson Immunoresearch, cat #111-035-003, 1:5000) for 1 hour at room temperature. As necessary, the blot was stripped using Restore PLUS Stripping Buffer (Thermo Fisher Scientific, Waltham, MA). The signal was detected using the iBright Imaging System, and densitometry of Western blot images was analyzed using ImageJ software and normalized to β-actin.

### Metabolic gene expression analysis using Nanostring nCounter system

Gene expression profiling in the knockout mouse model was performed using the NanoString nCounter Analysis System (NanoString Technologies, Seattle, WA, USA). Total RNA was extracted from the left ventricles of the heart using the RNeasy Fibrous Tissue Mini Kit (QIAGEN, cat No. 74704) according to the manufacturer’s instructions. The quality and concentration of RNA were assessed using a NanoDrop spectrophotometer. For the analysis, a NanoString nCounter Metabolic Pathway Panel was utilized, which included probes for 768 genes relevant to studying metabolism in the context of cancer, immunology, and metabolic diseases. The panel also included 20 internal reference genes for data normalization. Hybridization reactions were set up following NanoString protocols, with 20 μl of total RNA at concentrations ranging from 75 to 350 ng/µl per sample. The samples were processed on the nCounter Analysis System following the manufacturer’s instructions at the Iowa Institute of Human Genetics Core Facility. Data were collected and initially processed using nSolver Analysis Software (NanoString Technologies). Genes with a fold change of ≥1.25 and a p-value<0.05 were considered differentially expressed.

### mRNA sequencing

Total RNA was extracted from the left ventricles of six mice in each group using the RNeasy Fibrous Tissue Mini Kit (QIAGEN, cat No. 74704). All left ventricles of mice were collected in one day, and all RNA samples were prepared in one day. RNA integrity was assessed with the Agilent 2100 Bioanalyzer (Agilent Technologies), and samples with an RNA integrity number (RIN) ≥ 8 were sequenced on the Illumina NovaSeq 6000 platform at the University of Iowa Institute of Human Genetics Genomics Division. Libraries were prepared using the TruSeq RNA Library Prep Kit v2 (Illumina). Quality control of RNA-seq reads was performed using FastQC (version 0.1.2) and Trim Galore! (Version 0.6.5, Babraham Bioinformatics). The cleaned reads were aligned to the mouse reference genome (GRCm38) and quantified using Salmon. DESeq2 was used for normalization and differential gene-expression analysis. Genes with a Benjamini–Hochberg-adjusted P value (false-discovery rate, FDR) below 0.05 and an absolute log_2_fold change greater than 1 were considered differentially expressed. Principal component analysis was performed using Ingenuity Pathway Analysis (IPA; Qiagen). Gene set enrichment analysis was performed using GSEAPreranked in GSEA v.4.4.0 (Broad Institute) against the MSigDB Hallmark and KEGG gene set collections. Volcano plots, heat maps were generated using Python v.3.11 with matplotlib v.3.8. For heat-map visualization, expression values were standardized by row as z-scores and clipped at ±2.5. Genes within each pathway group were hierarchically clustered using average linkage and Euclidean distance. Joint pathway analysis integrating transcriptomic and metabolomic datasets was performed using MetaboAnalyst v.5.0. Genes associated with cardiac hypertrophy and remodelling were visualized using log_2_fold-change values obtained from DESeq2, with an FDR below 0.05 considered statistically significant.

### Metabolomic profiling by LC-MS

Mice were anesthetized as described above. After achieving deep anesthesia, rapid thoracotomy was performed, and the heart was excised. To preserve metabolite integrity, the heart was immediately flash-frozen in liquid nitrogen-cooled aluminum tongs, ensuring that the time from thoracotomy to freezing did not exceed 6 seconds. All the tissue samples were collected in one day. Metabolomic profiling was conducted at the University of Iowa Metabolomics Core Facility. Lyophilized tissue samples were homogenized, centrifuged, and the supernatants were dried. Dried extracts were reconstituted in acetonitrile/water (1:1 v/v) for LC-MS analysis on a Thermo Q Exactive hybrid quadrupole Orbitrap mass spectrometer coupled with a Vanquish Flex or Horizon UHPLC system. Chromatographic separation was achieved on a Millipore SeQuant ZIC-pHILIC column with a gradient of 20 mM ammonium carbonate and acetonitrile. The mass spectrometer operated in full-scan, polarity-switching mode with a scan range of m/z 70–1,000. Data were processed using Thermo Scientific TraceFinder 5.2 software, metabolites were identified based on the core facility’s in-house library, signal drift was corrected using NOREVA ^58^, and data were normalized to the sum of all measured metabolite ions. Statistical analysis of the metabolomic data was conducted using MetaboAnalyst to identify significantly altered metabolites between experimental groups.

### *In vivo* ^13^C isotope tracing in myocardium

On the day of the experiment, mice were subjected to 4 hours of food restriction and then received an intraperitoneal injection of either ^13^C-labeled (C6, Cambridge Isotope Laboratories, Tewksbury, MA) or unlabeled D-glucose at 1 mg/g body weight dissolved in 100 µl sterile saline. After 20 minutes, mice were euthanized under isoflurane gas, and hearts rapidly excised and frozen in liquid nitrogen cooled aluminum tongs as described above. The ^13^C isotope tracing analysis of substrates within the metabolite pools were analyzed and calculated as previously described ^59, 60^. Briefly, tissue samples were lyophilized and transferred to homogenization tubes. Extraction solvent consisting of acetonitrile:methanol:water (2:2:1, v/v/v) was added at 18× the tissue amount. The extraction solvent was spiked with heavy isotope-labeled internal standards (D8-Valine, ^13^C4, D3, ^15^N-Aspartate, ^13^C4, D4-Succinate) prior to sample homogenization. Samples were homogenized, rotated at −20°C for 1 hour, and centrifuged at high speed for 10 minutes. Aliquots of 100ul supernatant was transferred to clean 1.5 mL Eppendorf tubes and dried using a SpeedVac concentrator. Dried extracts were reconstituted in 50 µL of acetonitrile:water (1:1, v/v), vortexed thoroughly, and incubated at −20°C overnight to precipitate residual proteins. Samples were centrifuged again at high speed, and the resulting supernatants were transferred to LC-MS vials for analysis. LC-MS data were acquired as described in previous section, and inclusion lists were generated for all the metabolites and heavy internal standards in both positive and negative modes. Isolation window was set at 12 m/z and isolation offset was 5 m/z. Resolution was 70,000. During analysis, natural abundance samples (samples that are not treated with isotopic metabolite) were used for natural abundance correction.

### Statistical analysis

Statistical analyses were performed using GraphPad Prism v.9. All statistical tests were two-sided unless otherwise stated. Data are presented as mean ± s.d., and individual data points represent independent biological replicates. Sample sizes and the definition of n are provided in the corresponding figure legends. Comparisons between two independent groups were performed using an unpaired two-tailed Student’s t test. Comparisons involving three or more groups or multiple experimental factors were performed using one- or two-way ANOVA, as appropriate, followed by Tukey’s or Šidák’s multiple-comparisons test, as specified in the corresponding figure legends. Longitudinal echocardiographic measurements were analyzed using two-way repeated-measures ANOVA followed by Šidák’s multiple-comparisons test. Survival curves were analyzed using the Kaplan–Meier method and compared using the log-rank (Mantel–Cox) test. For all analyses, p<0.05 was considered statistically significant.

## Notes

### Competing Interest Statement

The authors have declared no competing interest.

