## supplemental figures for "Cardiomyocyte prohibitin ablation reprograms cardiac metabolism revealing a pathogenic role for mTORC1 in dilated cardiomyopathy"

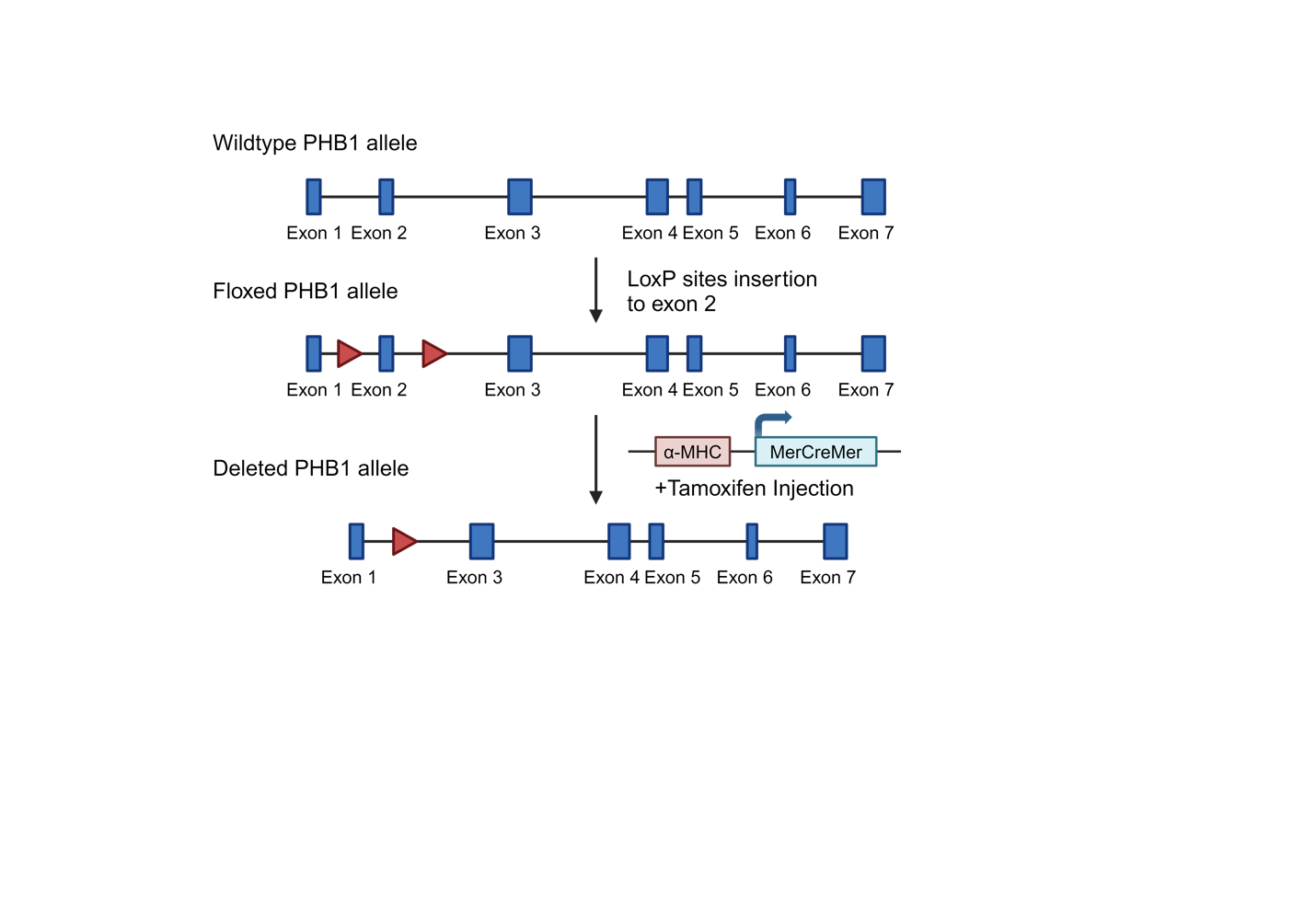

**Extended Data Figure 1: Schematic of the conditional PHB1 knockout strategy using the tamoxifen-inducible α-MHC-MerCreMer system.**

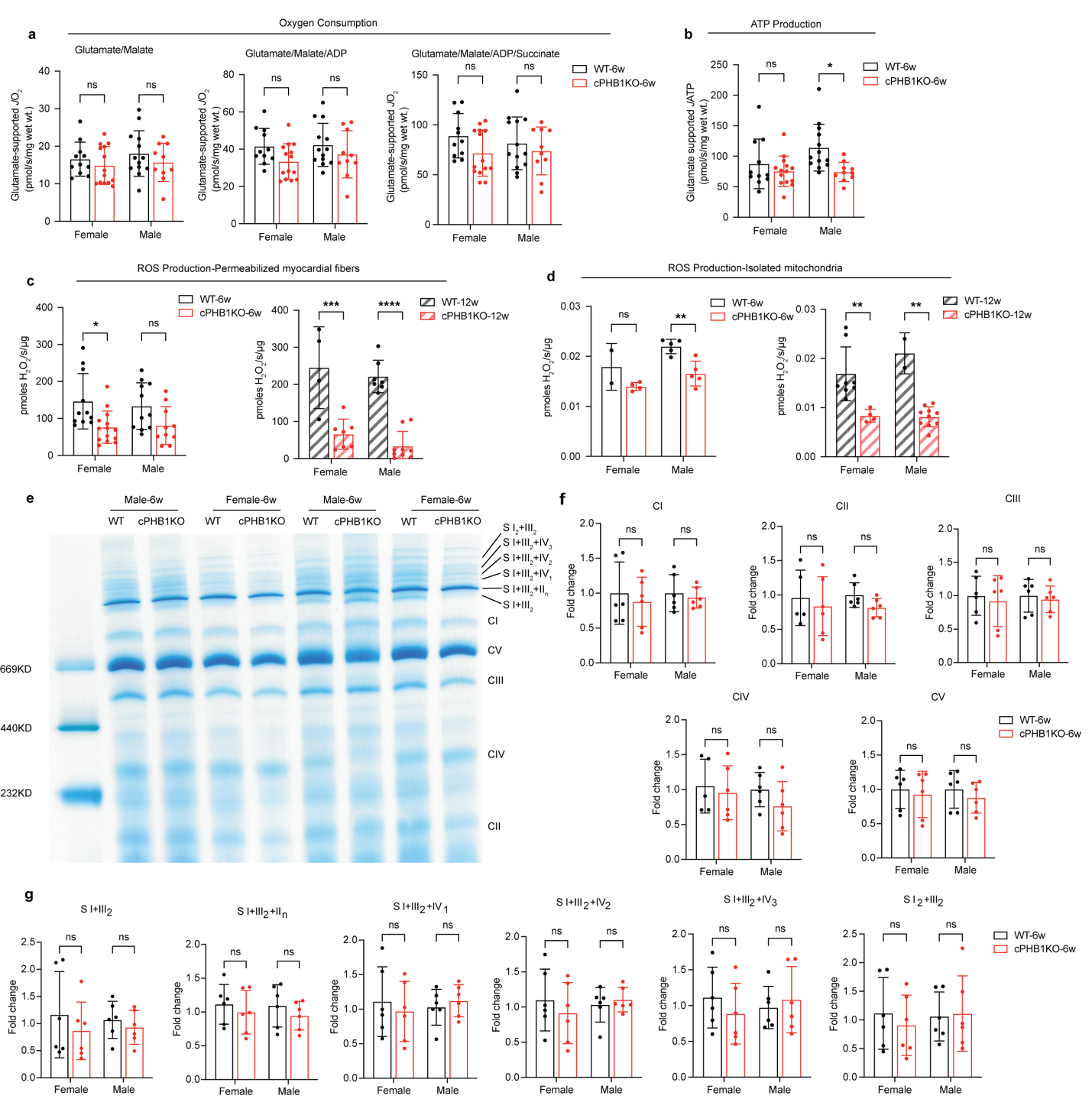

**Extended Data Figure 2: Mitochondrial respiration, ROS and respiratory supercomplex formation in cPHB1KO hearts.**

**a**,**b**, Mitochondrial respiratory flux (JO₂) (**a**) and ATP production (JATP) (**b**) supported by glutamate in permeabilized myocardial fibers from mice at 6 weeks post-tamoxifen. Data are mean ± s.d. Female: WT-6w, *n* = 11; cPHB1KO-6w, *n* = 14. Male: WT-6w, *n* = 13; cPHB1KO-6w, *n* = 10 mice. Significance was determined by two-way ANOVA with Tukey's multiple-comparisons test. **c**,**d**, H₂O₂ production measured by the reverse electron transport (RET) method with excess succinate in permeabilized myocardial fibers (**c**) and in isolated mitochondria (**d**) from female and male WT and cPHB1KO mice at 6 weeks (6w) and 12 weeks (12w) post-tamoxifen. Data are mean ± s.d. Significance was determined by two-way ANOVA with Tukey's multiple-comparisons test. **e**, Representative blue native PAGE gel with the indicated bands. Gels/blots were processed in parallel. Data are representative of two biological replicates per group. **f**,**g**, Quantification of individual mitochondrial complexes (**f**) and of supercomplex assemblies (**g**). Band intensities were normalized to the mean of the WT group within each sex and expressed as fold change, so each WT sample retains its spread around 1.0. Data are mean ± s.d., *n* = 6 biological replicates per group pooled from three gels. Statistical significance was determined by two-way ANOVA with Tukey's multiple-comparisons test. For all panels, *P < 0.05, **P < 0.01, ***P < 0.001, ****P < 0.0001. n.s., not significant.

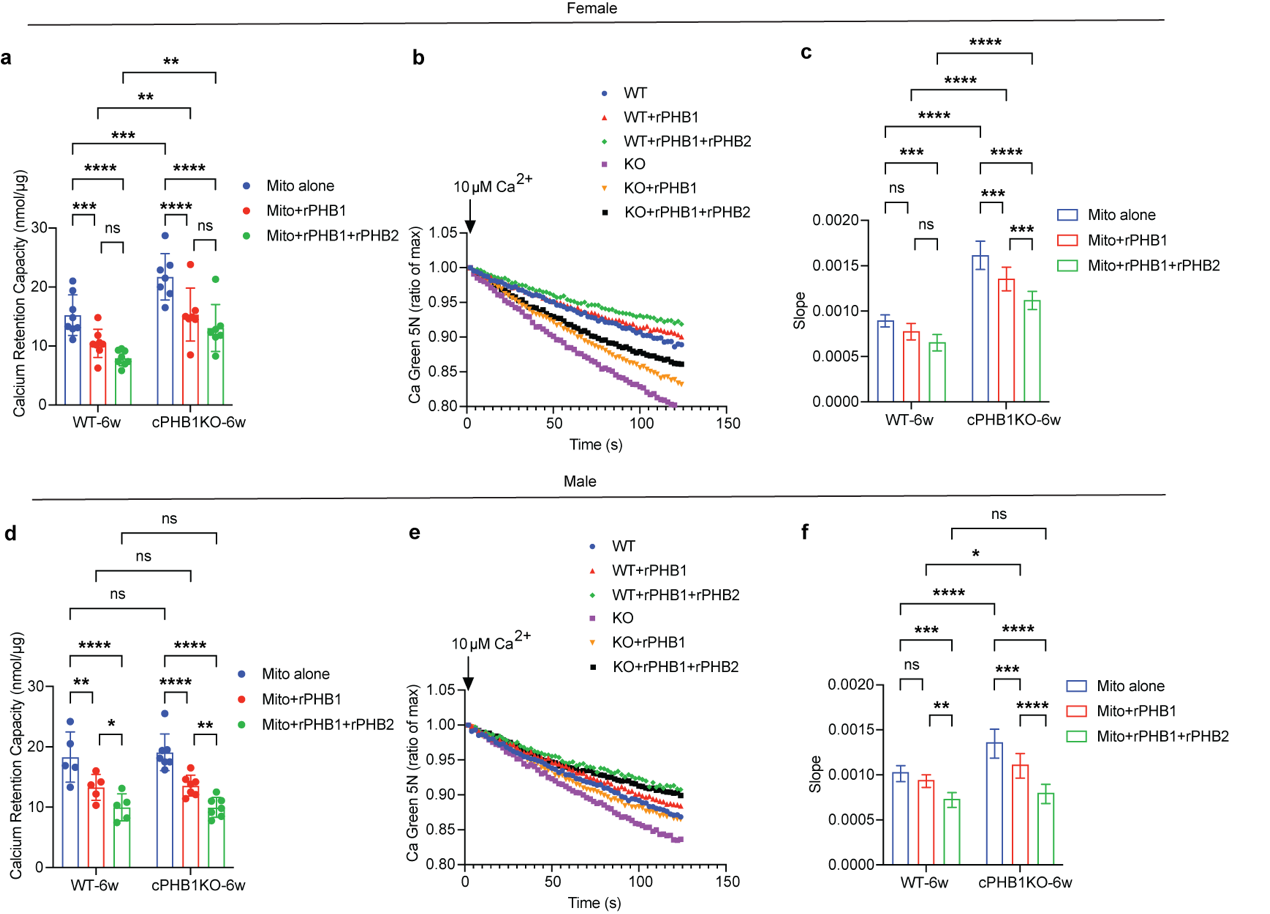

**Extended Data Figure 3: Recombinant PHB1 and PHB2 restores mitochondrial Ca^2+^ handling in cPHB1KO mitochondria.**

**a-c**, Female. **d**-**f**, Male. **a**,**d**, Quantification of calcium retention capacity in WT and cPHB1KO mice at 6 weeks post-tamoxifen. **b**,**e**, Representative traces of the second calcium-uptake peak from **a** and **d**. **c**,**f**, Slope of the linear regression fitted to the descending phase of the second fluorescence peak in **b** and **e**. Data are mean ± s.d. Female: WT-6w, *n* = 8; cPHB1KO-6w, *n* = 7. Male: WT-6w, *n* = 5 (calcium retention capacity) or 6 (slope); cPHB1KO-6w, *n* = 7 mice. For bar graphs, statistical significance was determined by two-way ANOVA with Tukey's multiple-comparisons test. For all panels, *P < 0.05, **P < 0.01, ***P < 0.001, ****P < 0.0001. n.s., not significant.

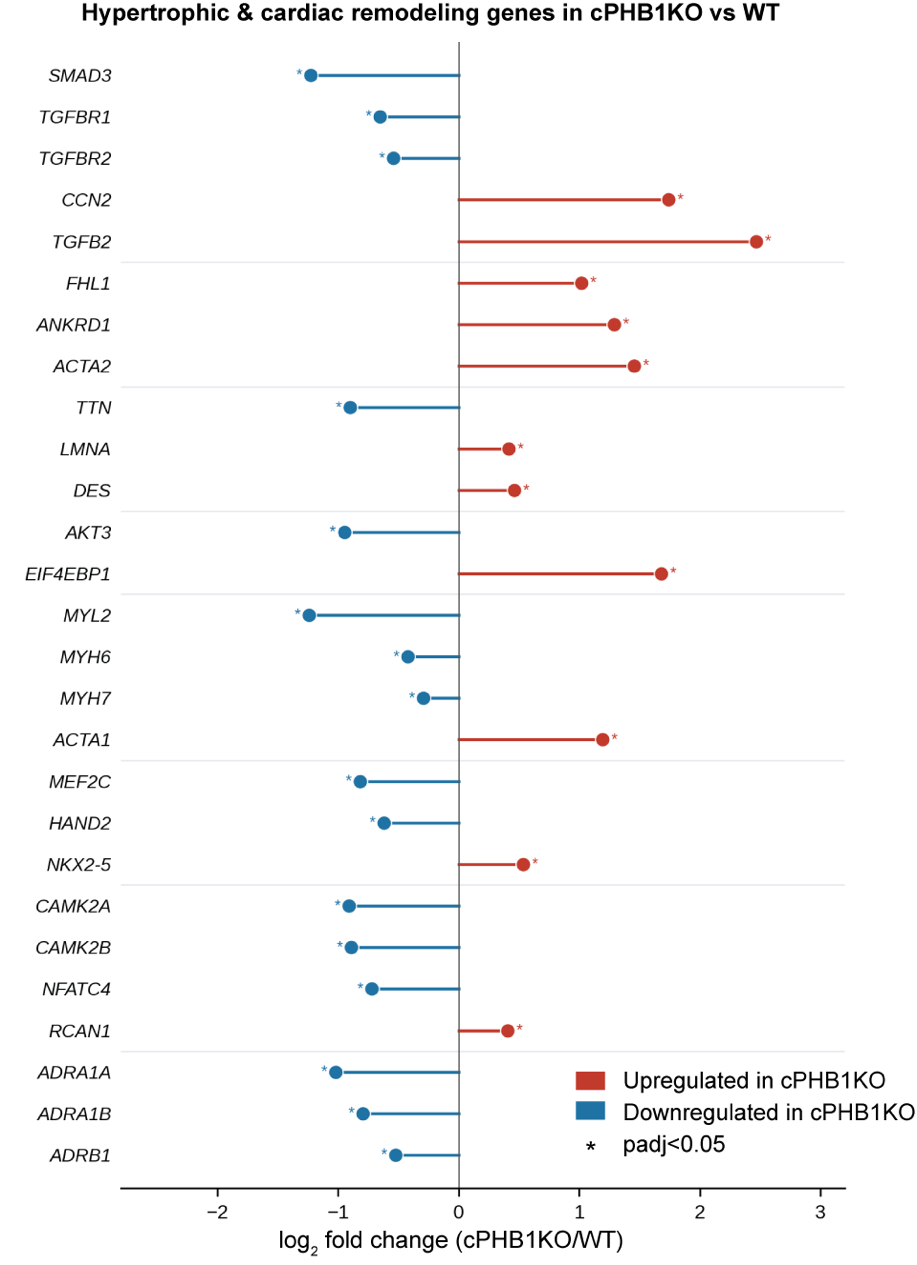

**Extended Data Fig. 4: Changes in hypertrophic and cardiac remodeling genes in cPHB1KO hearts.**

Log₂ fold change of hypertrophic and cardiac remodeling genes in cPHB1KO versus WT left ventricle from mRNA sequencing at 6 weeks post-tamoxifen. Genes upregulated in cPHB1KO are shown in red and downregulated genes in blue. All genes shown were significantly differentially expressed (DESeq2, adjusted P < 0.05). n = 12 per group (WT and cPHB1KO), 6 females and 6 males

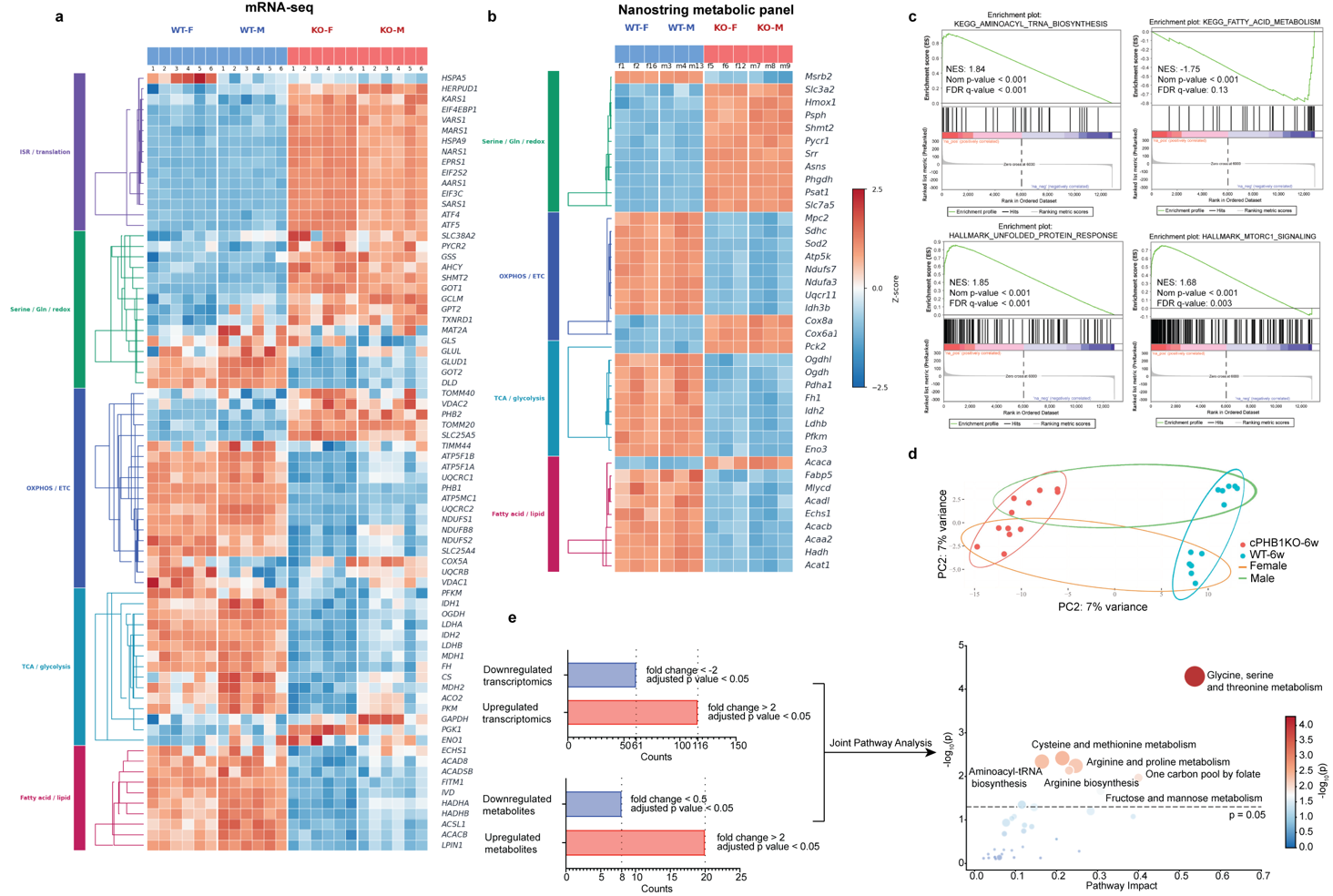

**Extended Data Fig. 5: Transcriptomic and metabolic pathway analysis of cPHB1KO hearts.**

**a,b**, Heat maps of differentially expressed genes from mRNA sequencing (**a**) and the NanoString metabolic panel (**b**), grouped by the pathways indicated on the left. mRNA sequencing: n = 12 per group (WT and cPHB1KO), 6 females and 6 males. NanoString metabolic profiling: n = 6 per group (WT and cPHB1KO), 3 females and 3 males. **c**, Gene set enrichment analysis (GSEA) of mRNA sequencing data for selected pathways using GSEA (Broad Institute, Subramanian et al., 2005) against the MSigDB Hallmark and KEGG gene set collections. **d**, Principal component analysis (PCA) of mRNA sequencing data with distinct gene expression profiles between genotypes. n = 6 per group (WT and cPHB1KO). **e**, Integrated joint pathway analysis (JPA) combining significantly changed genes (fold change >2 or <−2) from mRNA sequencing with metabolites (fold change >2 or <0.5) from metabolomics. Bubble size and position reflect pathway impact score (the ratio of observed to total genes in each pathway) and adjusted P value.

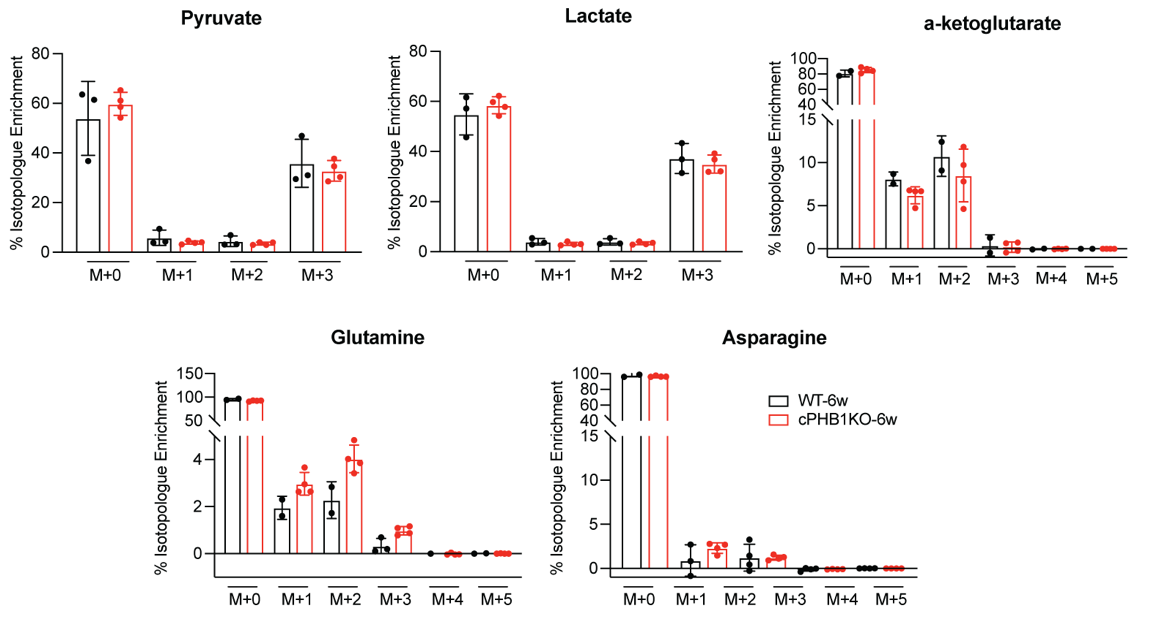

**Extended Data Fig. 6: Quantification of additional ¹³C isotope tracing.**

Data are mean ± s.d. WT-6w, *n* = 2 or 3; cPHB1KO-6w, *n* = 4 mice.

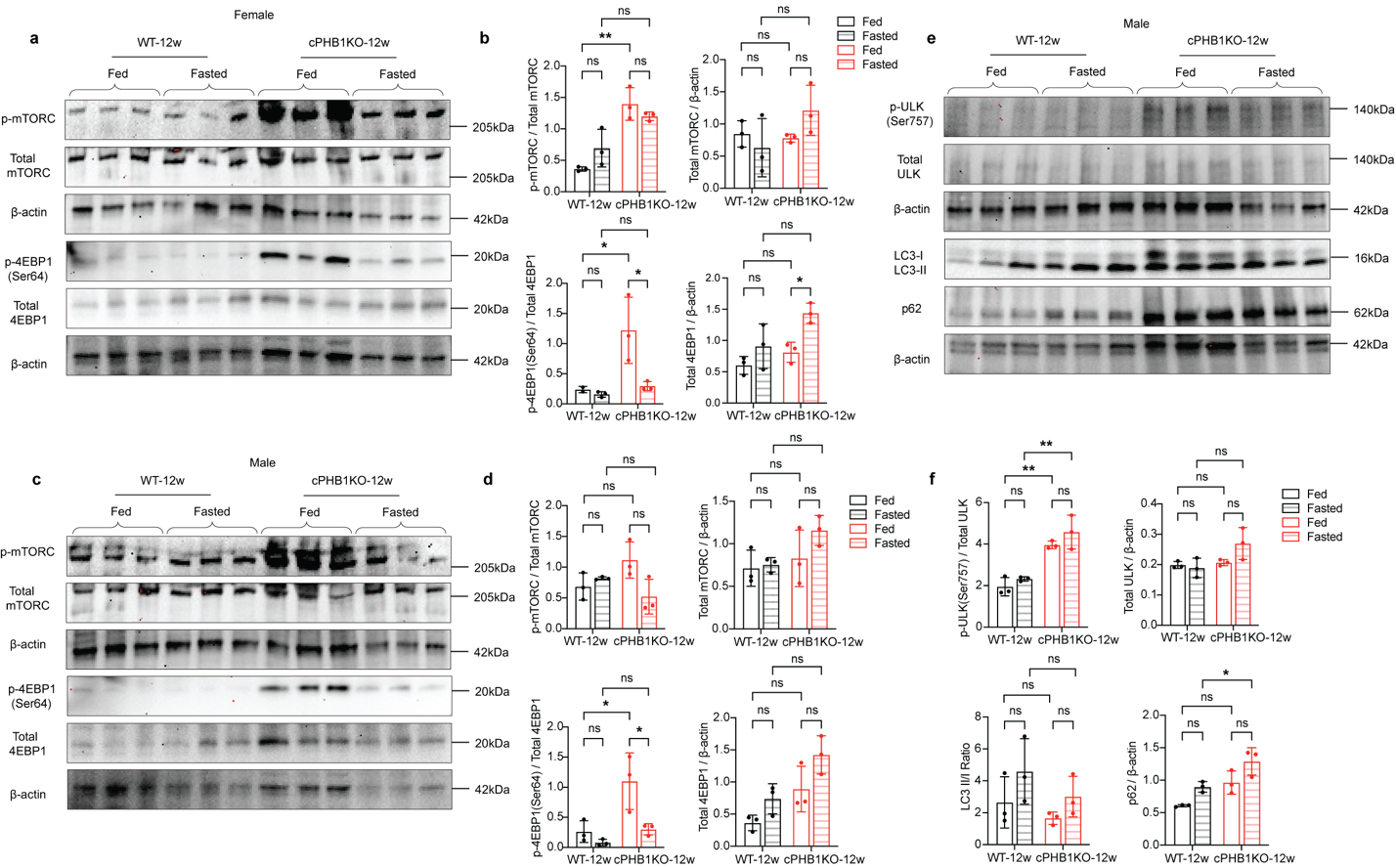

**Extended Data Figure 7: Effects of prolonged fasting on mTOR signaling and autophagy in WT and cPHB1KO hearts.**

**a**,**c**, Western blot of mTOR-pathway proteins phospho-mTOR (p-mTOR), total mTOR, phospho-4EBP1 (Ser64) (p-4EBP1 Ser64) and total 4EBP1 in left ventricular tissue extracts from female (**a**) and male (**c**) WT and cPHB1KO mice at 12 weeks post-tamoxifen (WT-12w and cPHB1KO-12w) under fed and fasted conditions. **Gels/blots were processed in parallel. Data are representative of three biological replicates.** β-actin was used as the loading control. **b**,**d**, Quantification of the immunoblots in **a** and **c**, for female (**b**) and male (**d**) mice, respectively. Data are mean ± s.d. *n* = 3 per group. Significance was determined by two-way ANOVA with Tukey’s multiple-comparisons test. **e**, Western blot of autophagy markers phospho-ULK1 (Ser757) (p-ULK Ser757), total ULK1, LC3-I and LC3-II, and p62 in left ventricle from male WT and cPHB1KO mice at 12 weeks post-tamoxifen under fed and fasted conditions. **Gels/blots were processed in parallel. Data are representative of three biological replicates.** β-actin was used as the loading control. **f**, Quantification of the immunoblots in **e**. Data are mean ± s.d. *n* = 3 per group. Statistical significance was determined by two-way ANOVA with Tukey’s multiple-comparisons test. For all panels, *P < 0.05, **P < 0.01. n.s., not significant.

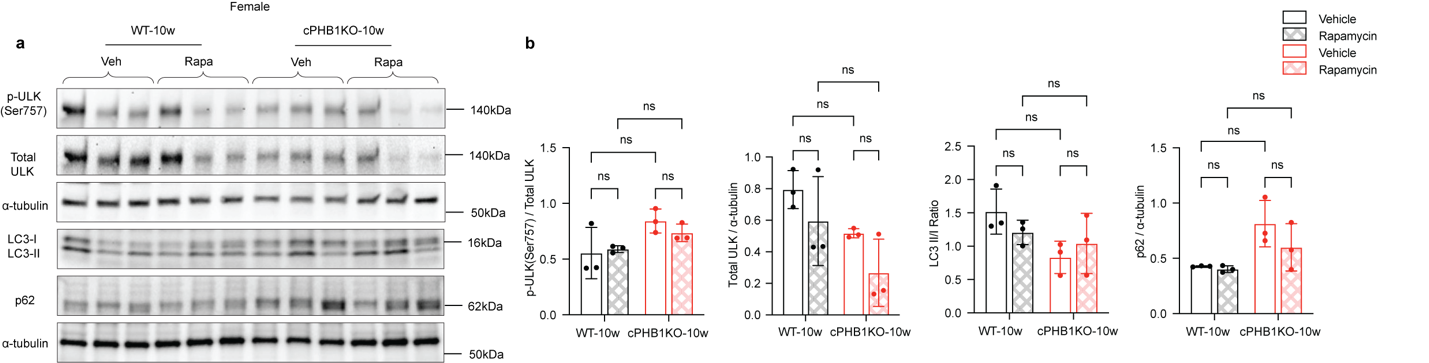

**Extended Data Figure 8: Rapamycin does not alter autophagy markers in female cPHB1KO hearts.**

**a**, Western blot of autophagy markers phospho-ULK1 (Ser757) (p-ULK Ser757), total ULK1, LC3-I and LC3-II, and p62 in left ventricular tissue extracts from female WT and cPHB1KO mice at 10 weeks post-tamoxifen (WT-10w and cPHB1KO-10w), treated with vehicle (Veh) or rapamycin (Rapa). Gels/blots were processed in parallel. Data are representative of three biological replicates. α-tubulin was used as the loading control. **b**, Quantification of the immunoblots in **a**. Data are mean ± s.d. *n* = 3 per group. Statistical significance was determined by two-way ANOVA with Tukey's multiple-comparisons test. n.s., not significant.
